# RNA binding protein Musashi1 interacts with the viral genomic RNA and restricts SARS-CoV-2 infection by repressing translation

**DOI:** 10.1101/2024.09.29.615653

**Authors:** Sourav Ganguli, Divya Gupta, Rajashree Ramaswamy, Rajashekar Varma Kadumuri, Aswathy G Krishnan, Dixit Tandel, Deena T David, Soumya Bunk, Sreenivas Chavali, Krishnan Harinivas Harshan, Pavithra L. Chavali

## Abstract

Musashi RNA binding proteins are important post-transcriptional regulators of stem cell homeostasis and are known to be involved in viral infections. However, their role in SARS-CoV-2 infection remains largely unknown. Using computational studies, *in vivo* RNA immunoprecipitation and biochemical assays, here, we establish that Musashi1 (Msi1) interacts with viral genomic RNA through direct binding to the SARS-CoV-2 3’UTR. Importantly, binding of Msi1 to the viral 3’UTR results in translational repression mediated by inhibition of Poly (A) binding protein (PABP). Conversely, Msi1 knockout promotes robust viral replication and increased viral protein expression. Using 2D cell cultures, stem cells and 3D organoids, we show that depletion of Msi1 in intestinal cells augments infection. This finding explains why the human intestine serves as a reservoir for the SARS-CoV-2 virus, wherein differentiated enterocytes, which have negligible levels of Msi1, are highly affected. Contrarily, stem cells which are enriched for Msi1 expression, are known to be less permissive to SARS-CoV-2 infection despite expressing the entry receptors. Our findings show how translation repression of SARS-CoV-2 by stem cell RNA binding proteins such as Msi1 could help evasion of infection.

## INTRODUCTION

RNA viruses constitute the major fraction of the transmissible emerging pathogens, comprising of about 223 geocoded viruses (1). Since RNA viruses cause several infections and lead to high mortality, understanding molecular mechanisms underlying their pathogenesis is paramount to develop effective intervention strategies. In this context, the role of host RNA binding proteins (RBPs) which interact with the viral RNA genome and modulate viral life cycle assumes significance to understand viral biology and design efficient inhibitors. Host RBPs are important for viral biology due to the dependence of the viral RNA genome on host cellular machinery for translation. Thus the host RBPs predominantly affect the outcome of viral infection by affecting viral replication, stability and translation (2). For instance, many flaviviruses are enriched for Musashi binding elements, which are recognized and bound by post transcriptional regulators Musashi1 (Msi1) and Musashi2 (Msi2) (3–5). We previously showed that Msi1, a stem cell RBP, binds to the Zika RNA and promotes its replication by specifically binding to the 3’UTR of the viral genome (3). Interestingly, overexpression of Msi1 in Zika non-permissive HEK293 cells promoted viral replication demonstrating how RBPs could influence cellular/tissue landscape of viral infection. Notably, RNA viruses are adept at molecular mimicry, exemplified by the rapid acquisition of RBP binding elements that sponge cellular RBPs often resulting in beneficial outcomes for the viruses. For example, the more infective Brazilian strain of Zika had acquired an additional Msi binding element (GUAG) in its 3’UTR and showed an increased binding to the protein. This enables Zika viral RNAs to compete with the host RNAs for Msi1 binding resulting in additional viral burden in the neural progenitor cells (3).

Conserved RNA binding domains (RBDs) encoded within RBPs enable sequence-dependent recognition of an RNA. These RBDs typically recognize relatively short nucleotide sequences or preferentially bind to more simplified nucleotide patterns, such as purine-rich or pyrimidine-rich tracts of RNA. For instance, Msi1 and Msi2 bind RNA consensus sequences: 5’-[GA]U(1–3)AGU-3’, through two conserved tandem RNA recognition motifs (RRMs; RRM1 and RRM2) (6). In addition to the RRM sequences and cognate sites, the tissue context might also play a role in viral infection. For example, fetal neural progenitors are the prime targets of Zika virus, while SARS-CoV-2 primarily infect differentiated cells (7–9). SARS-CoV-2 infects the respiratory tract via angiotensin-converting enzyme 2 (ACE2) receptor attachment (10). Once the virus enters pulmonary site by engaging its Spike protein with ACE2 or the transmembrane serine protease protein 2 (TMPRSS2), it can be either cleared from the mucociliary tract or can enter gastrointestinal (GI) tract. The GI tract can serve as a reservoir for SARS-CoV-2, since the fecal samples of about 48%-54% of COVID-19 patients were positive for viral RNA, and about 15%-17% of patients exhibited gastrointestinal symptoms (11–14). The virus can then trigger a cytokine storm leading to moderate to severe pathologic manifestations (15,16).

The genomic RNA (gRNA) of SARS-CoV-2 serves as a bifunctional template, facilitating the synthesis of both full-length negative-sense RNAs (ncrRNAs) required for genome replication and various sub-genomic negative-sense RNAs (sgRNAs) that are subsequently translated into corresponding sub-genomic mRNAs (17). Akin to other RNA viruses, SARS-CoV-2 depends on host proteins for the assembly of its replication and translation machinery (18). Several studies have thoroughly characterized the SARS-CoV-2 RNA-protein interactome (19–22). These studies show that the negative-sense ncrRNAs are associated with an exceptionally diverse and extensive array of host proteins, which are involved in regulating viral production, apoptotic signaling, and immune responses. This underscores the critical role that negative-sense RNAs play during infection. For instance, the viral 5’UTR interacts with host proteins from the U1 small nuclear ribonucleoprotein (snRNP) family, while the viral 3’UTR interacts with host proteins associated with stress granules and heterogeneous nuclear ribonucleoprotein (hnRNP) families. (23). Notably, the conserved elements in 3’UTR of the SARS-CoV-2 comprises several domains that are important for controlling viral RNA production (24). Here, we report that SARS-CoV-2 gRNA contains conserved Musashi binding elements (MBEs) in its 3’UTR. We further show that Msi1 binds to the MBEs in the viral 3’UTR and this interaction limits the viral load, by repressing the translation of the virus.

## MATERIALS AND METHODS

### Estimation of binding sites opening energies based on single strandedness

Following the method previously described (5), we employed a biophysical model that characterizes RNA at the secondary structure level, building upon the thermodynamic nearest neighbour energy model from the ViennaRNA Package (25). This model facilitates the computation of RNA’s most stable, minimum free energy (MFE) structure and the partition function, *Z*. The equilibrium probability of a secondary structure, is determined by the equation *p*(*s*)=(*e*−*E*(*s*)/*RT*)/*Z*. Efficiently computing the partition function, individual base pair probabilities can be determined for extensive sequences. The accessibility, or the probability that a segment along the RNA remains single-stranded, derives from this partition function. The model’s accessibility (probability of an RNA segment being single-stranded) is derived from this partition function. To this RNAplfold from the ViennaRNA Package was used to compute the local pairing probabilities of GUAG, AUAG and AGAA tetranucleotide motifs to gauge the single-strandedness.

We assessed the significance of the binding element opening energy by comparing it with a total of 1000 randomized sequences, maintaining the dinucleotide composition. Opening energies in both genomic and shuffled sequence contexts were compared using z score statistics, defined as Z=*E*open(*WXYZ*)−*μ*/*σ*. Where *E*open(*WXYZ*) is the opening energy of tetra nucleotide *WXYZ* in its genomic context, while *μ* and *σ* signify the mean and standard deviation, respectively, of the opening energies for *WXYZ*, determined from an extensive set of randomized sequences. Randomization involved dinucleotide shuffling of the 100 nucleotide windows upstream and downstream of binding site, keeping tetra nucleotide motif sequence in place. To this inhouse python script is developed encapsulating the methodology provided in Perl utility plfoldz.pl, available at https://github.com/mtw/plfoldz. The tool employs the ViennaRNA for thermodynamics calculations, reporting tetranucleotide opening energies alongside a z score from 1000 dinucleotide shuffling events.

### Molecular Dynamics simulations

All-atom molecular dynamics simulations were performed using the GROMACS 2019.3 simulation package (26,27). The Charm36 force field, tailored for the protein–RNA complex, was sourced from a prior study (28). This complex was solvated in a dodecahedron box using the TIP3P water model, ensuring a minimum distance cutoff of 10 Å between the complex’s surface and the box edge. The system was neutralized with Na^+^ ions, and energy minimization was achieved using the steepest-descent algorithm until the energy converged to less than 1000 kJ/mol/nm. Before production run, all systems were equilibrated for pressure and temperature by position restraining the protein–RNA for 100 ps under both NPT (Number of particles, Pressure, Temperature) and NVT (Number of particles, Volume, Temperature) ensembles at designated temperatures. Long-range electrostatic interactions were computed using the Particle Mesh Ewald (PME) method (29) with a 0.16 nm grid spacing. A cutoff of 1.0 nm was designated for short-range electrostatic and van der Waals interactions. The LINCS algorithm (30) was used to constrain the bond lengths. All simulations were performed for a simulation timescale of 50 nanoseconds at 300 Kelvin temperature using a 2-fs integration time step, with a coupling coefficient of tT=0.1 ps using modified Berendsen thermostat (31), and Parrinello–Rahman pressure-coupling at 1 bar with a coupling coefficient of tP=1 ps. Finally Data analysis was conducted using the Gromacs 2019.3 package.

### Computational design of RNA 2D structure and Protein-RNA complex

The secondary structure of the reference 3’UTR sequence, possessing the lowest free energy, was predicted using the RNAstructure Fold webserver (32–34). The analysis was conducted under default parameters at a temperature of 310.15 Kelvin. Final conformation with a probability greater than 99% and a free energy of –74.7 kcal/mol was chosen for subsequent analysis. The two-dimensional representation of the predicted RNA structure was generated using the RNA2Drawer webserver (35). To investigate the binding efficiency Zika virus 3’UTR (WT) with the two MBEs in SARS-CoV-2, M1(SL1) and M2 (HVR1) were modelled using Discovery Studio Visualizer. Assisted docking of Human Musashi 1 (PDBID:5X3Z) with the designed RNA substrates were modelled in PyMOL, with inference from the solution structure of Msi1 RNA binding domain 2 in complex with RNA (PDBID:5X3Z).

### Cell Culture

Caco-2 cells were maintained in DMEM GlutaMax (Gibco), supplemented with 20% (v/v) Fetal Bovine Serum (FBS - Gibco). HEK293T cells were maintained in DMEM GlutaMax (Gibco), supplemented with 10% (v/v) Fetal Bovine Serum (FBS - Gibco). Induced pluripotent stem cells were purchased from ThermoFisher and maintained in mTESR media. All the cells were grown and incubated at 37°C and 5% CO2 and were tested every month for mycoplasma using a PCR based assay. Intestinal organoids were generated using Stemdiff intestinal organoid kit (Stem Cell Technologies Inc) and endoderm identity confirmed with SOX17 staining.

### Constructs and Primers

PUC57 vector containing SARS-CoV-2 minigene with Gaussia luciferase sequence was obtained from Dr. Jingxin Wang (18). From this, both the 5’UTR and the 3’UTR were PCR amplified and cloned into pJet1.2 blunt vector. To generate the M1 and M2 mutants, megaprimer based mutagenesis (36) was performed using pJet1.2 3’UTR as the template with the following substitutions:

Stem loop 1 – GUAG >> GUGC at 113 nucleotide

Hypervariable Region 1 – GUAG >> GUGC at 292 nucleotide

Subsequently, double mutant was generated using a similar strategy, using M1mutant as the template. Using this method, SARS-CoV2 minigene construct was mutated to generate a double mutant minigene construct for Gaussia luciferase assay.

For the firefly luciferase assay, Wild type or the double mutant 3’UTR from pJET1.2 were cloned into pGL4.13 downstream of Firefly luciferase, by digestion with XbaI-HF(NEB). ShRNA targeting MSI1 and the scrambled shRNA were annealed using the established protocol and ligated to AgeI-HF (NEB) and EcoRI-HF (NEB) digested pLKO.1. DNA sequencing was used to confirm the sequence. Primers used in this study are listed in **Table 1**.

**Table 1.**
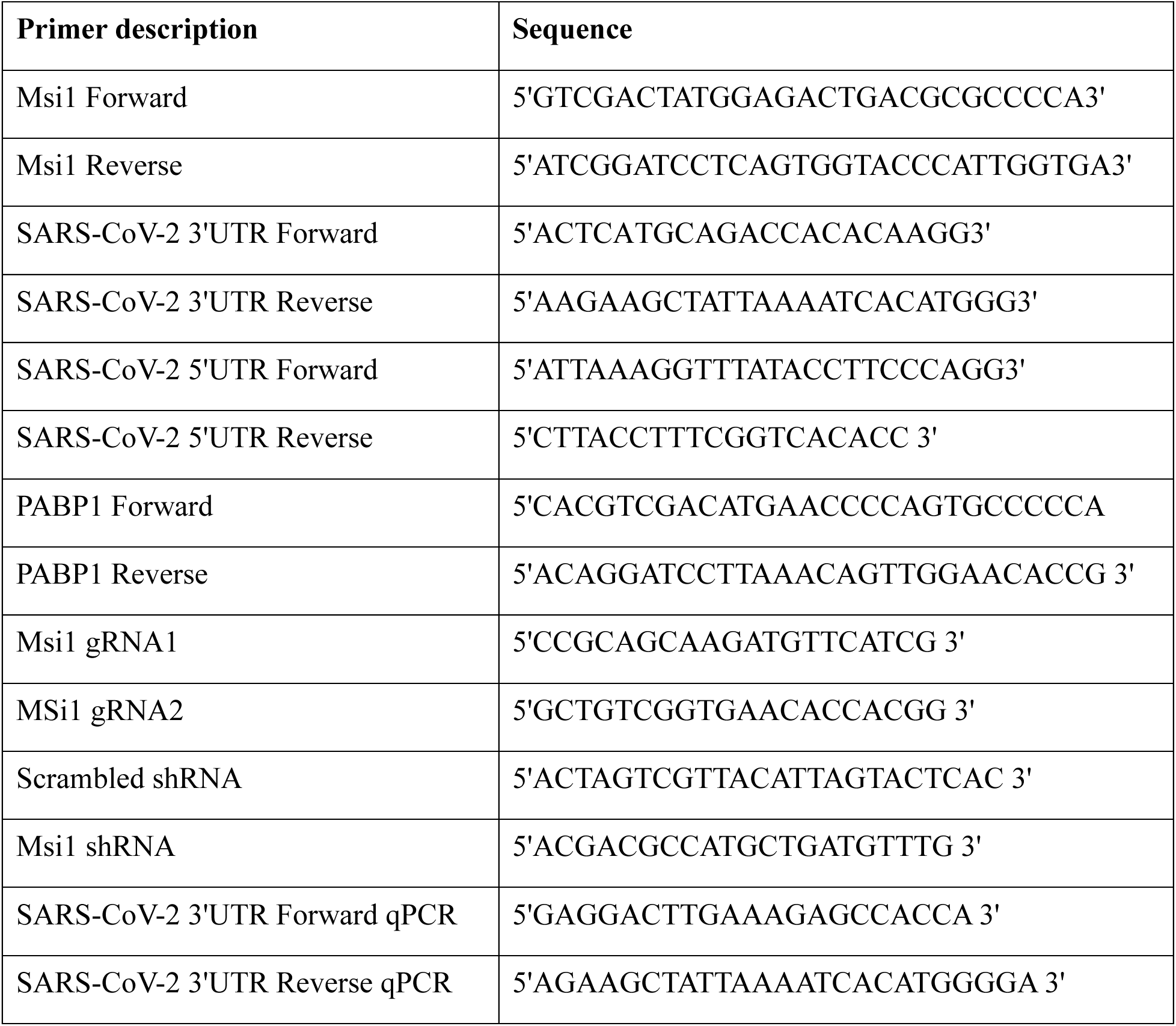
List of primers used in the study.

### Lentiviral Preparation and genomic modifications of cell lines

Cas9 lentivirus was generated by transfecting 10μg of Lenti-Cas9-Blasticidin plasmid (Transomics) in HEK293T cells along with 2.5μg PAX-2 and 7.5μg of PMD2. Caco-2 cells stably expressing Cas9 were generated after 10μg/ml of blasticidin selection for a week. Msi1 knock-outs were then generated in this background (Parental) using transEDIT-dual CRISPR gRNA lentiviral expression vector (TEDH-1049415) with target sequence of 5’CCGCAGCAAGATGTTCATCG3’ and 5’GCTGTCGGTGAACACCACGG3’. These cells were selected with puromycin (1.5μg/ml for 96 hours) to generate Musashi1 knockout-pool. Subsequently, they were diluted in a 96-well plate for clonal expansion of single genotypes and screened using western blotting to confirm the absence of Musashi1. Transient rescues were made using lipofectamine 3000 (Thermofisher) based expression of transgene Msi1-tagged with GFP.

### SARS-CoV-2 virus generation and infection

All viral experiments were carried out in the BSL-3 facility. Studies were performed using the B.1.1.8 strain of SARS-CoV-2, which was produced according to the methodology previously mentioned (37). Viral transport medium (VTM) having less than 20 CT value in real time quantification was filter sterilized (0.22μm) and utilized as inoculum for infecting Vero cells in a 96-well format. This cell culture supernatant was utilized to infect fresh Vero cells, after the cytopathic effect was seen. This procedure was repeated until the supernatant showed an infectious titre of 10^-7^ ml^-1^. For strain identification and confirmation, the viral culture was sequenced using next generation sequencing. Caco-2 Cells (Parental, Cas9 control and Musashi1 knockouts) were infected with MOI:1 (multiplicity of infection) in serum free DMEM for 3 h at 37°C with 5% CO2. Following which the cells were supplied with 20% serum containing DMEM. At 48 hours post infection (hpi) supernatant was collected for real time quantification and plaque assay, and cells were harvested for western blotting.

### RNA Immunoprecipitation assay

RNA Immunoprecipitation (RIP) was performed from the Caco-2 cells mock/infected with SARS-CoV-2 using rabbit IgG (Sigma) and MSI1 (ab21628) antibodies as previously reported (3). Quality of immunoprecipitations was assessed by immunoblot analysis. Atleast 1.5mg of protein lysate was made up to a volume of 500μl and incubated with 8μg of Musashi1 or IgG antibody for 3 h at 4°C, followed by addition of 50μl of prewashed protein A/G dynabeads (Life Technologies) for 2h. Beads were then washed 3 times for 3 min each with lysis buffer and RNA isolated after addition of 1 ml TRIZOL (Life Technologies cat. no. 15596-018). RNA isolated from 10% lysate was used as input. One step TB green mix (Takara) was then used for qPCR analysis to evaluate the binding to 3’UTR and Meril One-Step qRT-PCR Kit (Thermo Fisher) for E, and N binding.

### UV-Cross Linking Immunoprecipitation assay

UV-Cross Linking Immunoprecipitation (CLIP) was performed as described previously with minor modifications (3). Mock/Infected Caco-2 cells were crosslinked with 300mJ/cm2 of UVA (365nm) on ice using Stratalinker device. For the assay the pellets were resuspended in 1ml of NP40 lysis buffer and incubated for 15 mins with constant mixing every 2 min. The lysates were then centrifuged at 10,000g for 15 min at 4°C. Subsequently lysates were subjected to partial RNaseIF digestion (New England Biolabs, 5 μl 1:50 dilution per 1ml of cell lysate) for 5 min at 37°C and verified on agarose gel. This lysate was incubated with MSI1-coated or IgG Protein G Dynabeads (Life Technologies). Beads were then washed in NP40 lysis buffer and DNaseI (20 U, Promega) was added for 15 min at 37°C following addition of 0.1% SDS and 0.5 mg/ml Proteinase K (Thermo Fisher Scientific) for 15 min at 55°C. The supernatant was collected and RNA was extracted using phenol/chloroform extraction and ethanol precipitation. Finally, RNA was dissolved in 20 μl of RNase free water and qPCR was performed using One step TB green mix (Takara).

### SDS PAGE and western blotting

Whole cell lysates were loaded with equal amount of protein and resolved in a 10% SDS PAGE gel. The proteins were then transferred to a nitrocellulose membrane which was stained with ponceauS and imaged. After imaging the blot was incubated for 1h in 5% non-fat lyophilized milk dissolved in 1XTBST, or in 5%BSA made in 1XTBST. It was then incubated overnight at 4°C or 2h at room temperature with primary antibodies. The primary antibodies used are Anti-Musashi1 (1:1000, Cat# ab21628, Abcam); Nucleocapsid (1:8000, Cat# MA5-29981, Invitrogen in 5% BSA), Actin (1:10000, Cat# 3700-Cell Signaling technologies); Anti-Musashi2 (1:1000, Cat#ab76148, Abcam) ACE2 (1:1000, Cat#92845, Cell Signaling Technologies). After overnight incubation the blot was washed three times for 5 min each with 1xTBST to remove any non-specific binding of the primary antibody. The blot was then incubated for 1h at room temperature with the appropriate secondary antibody. The secondary antibodies commonly used in our study were anti-mouse HRP conjugated secondary (Cat# ab97023, Abcam 1:4000); Anti-Rabbit Secondary IgG (Cat# NB7115, Novus Bio,1:5000). After incubation the blot was again subjected to three washes and then developed using HRP-Luminol based chemiluminescence (Clarity, Biorad) in Vilber-Lourmat chemidoc machine. For densitometry, the observed bands were quantitated using ImageJ and normalized against the loading control.

### Real time quantification

RNA from cell culture supernatant was isolated using viral RNA isolation kit (MACHEREY-NAGEL GmbH & Co. KG). Real-time quantitative PCR (RT-PCR) was performed in Roche LightCycler 480 using commercial kits. LabGun™ COVID-19 RT-PCR Kit was used to measure the RNA levels of SARS-CoV-2 RdRp and E gene levels, following manufacturers’ protocol. Fold changes between samples were calculated by ΔΔCp method using the internal control (IC) for normalization The RNA was diluted with nuclease free water in a ratio of 1:40 and 1μl was used for a 10μl reaction with One Step CT Power SyBR green mix (Thermofisher) or TB green mix (Takara) as per the manufacturers’ instructions.

### Luciferase assay

HEK293T cells were plated in a 12well plate at 70% confluency and transfected with pGL4.13-3’ UTR along with empty pCDNA vector and pCDNA-Msi1 and pRL-SV40. After 24h, cells were lysed and luciferase assay performed using Dual Luciferase Reporter Assay System (Promega).

The data was obtained using a PerkinElmer Microplate reader and the ratio of firefly to renilla luciferase plotted.

For Gaussia luciferase assay, HEK293T cells were plated in 24 well plate at 70% confluency and transfected with empty peGFPC1 plasmid and peGFPc1-Msi1 plasmid. After 24 hours, equal amounts (145ng) of capped wild type or the mutant minigene RNA was transfected. Minigene RNA was generated using M13 forward and reverse PCR amplified fragment using mMessage mMachine T7 transcription kit as per manufacturer’s instructions. After 48 hours of minigene RNA transfection, cell culture media was collected and centrifuged at 1500g for 5mins to remove cell debris. 20 μl of the media was subjected to Luciferase assay using Gaussia glow assay kit (ThermoFisher).

### *In vitro* biotinylated RNA pulldown

Two micrograms of the wild type and mutant SARS-CoV-2 3’UTRs in pJet1.2 vector were linearised and subjected to *in vitro* transcription using Megashortscript T7 poymerase kit as per manufacturer’s instructions (Invitrogen). The mixture was incubated for 3h at 37°C and EDTA (60mM) was added and precipitated overnight at –20°C. After centrifugation at 14000g for 30 min, the precipitate was dissolved in 50μl of water and cleaned with Quick spin clean-up columns (Roche). Following the DNaseI treatment (Promega) for 15 min at 37°C the reaction was cleaned once again with Bio-Spin 30 columns and RNA was precipitated overnight at –20°C and dissolved in 30μl of RNase free water. Purity of RNA was analysed on agarose gel using RNA Gel Loading Dye (ThermoFisher Scientific) after denaturation at 72°C for 10 min.

For the pulldown, three hundred micrograms of total protein from Caco-2 cells were used (lysis buffer: 25mM Tris HCl, pH 7.4, 150mM KCl, 5mM EDTA, 5mM MgCl2, 1% NP40, 0.5mM DTT, protease inhibitor cocktail (Roche), 100U/ml RNAse OUT) and first precleared with Streptavidin beads (GenScript). The precleared lysate was diluted 2X in lysis buffer and supplemented with tRNA (0.1μg/μl, ThermoFisher Scientific) and incubated with different concentrations of biotinylated RNA (5-20 pmol) for 2h at 4°C. In case of pure protein pull down the appropriate amount of the Msi1 proteins were quantitated and used. Biotinylated RNA was heated up to 60°C for 5 min and slowly cooled down to room temperature. Subsequently, 50μl of Streptavidin beads were added for an hour and the beads were washed three times with lysis buffer containing 300mM KCl. Finally, the beads were resuspended in 50μl of sample buffer and boiled for 5 min at 95°C. After a short spin, the supernatant was collected and subjected to western blot analysis.

### Statistical analysis

Two-tailed t-tests were used to determine statistical significance of data in all experiments. P-values for each comparison are indicated in graphs. Each experiment was performed using a minimum of 3 biological replicates. All viral infections and qPCR experiments were performed double-blind.

## RESULTS

### The 3’UTR of SARS-CoV-2 genomic RNA contains conserved Musashi binding elements

Sequence analysis of the 5’UTR and 3’UTR across 224 variants of SARS-CoV-2 revealed the presence of two putative conserved Musashi binding elements (MBEs; defined by GU(1–3)AG) specifically in the 3’UTR (**Figure 1A, Supplementary Figure 1A**). The two putative MBEs (henceforth referred as M1 and M2) were located in the (i) conserved loop1 of Stem and Loop 1 structure (SL1; M1) and (ii) the more divergent hypervariable region (HVR; M2) (**Supplementary Figure 1A**). The SL1 structure is essential for viral replicability, while the HVR region may contribute to viral pathogenicity (38). Notably, both the MBEs are conserved in SARS-CoV and SARS-CoV-2, but not in Middle East respiratory syndrome (MERS), Bat or Mouse Hepatitis Virus (MHV) viruses (**Figure 1B**). We next estimated the free energy required to force the MBE to be single-stranded (opening energy) using a previously described method (5). Strikingly, across all variants including Omicron, we identified that GUAG in the MBEs exhibited low opening energies with Z scores ranging from –1.10 to 1.53, reflecting their accessibility in the genome throughout the evolution of SARS-CoV-2 (**Figure 1C, Supplementary Figure 1B**). Contrarily, both the AUAG and the recently shown AGAA motifs, which have the propensity to bind to Msi1 (6), showed a higher median distribution of binding energies, alluding to a possible unfavourable context for Msi1 binding (**Supplementary Figure 1C**). To examine if Msi1 binds to the SARS-CoV-2 3’UTR, we transfected GFP-tagged Msi1 into HEK293 cells and performed *in-vitro* RNA pull-down with biotinylated SARS-CoV-2 3’UTR, using Zika 3’UTR as a positive control. Indeed, we observed binding of Msi1 to both SARS-CoV-2 3’UTR and Zika 3’UTR (**Figure 1D**). To confirm that the endogenous Msi1 could bind to the 3’UTR of SARS-CoV-2, we performed RNA pull down with the colon carcinoma cell line (Caco-2) which expresses endogenous Msi proteins. We find that both Msi1 and Msi2 can bind to SARS-CoV-2 3’UTR (**Figure 1E**). These findings suggest that the 3’ UTR of SARS-CoV-2 contains well-conserved Musashi binding sites and is amenable to binding to Msi proteins.

**Figure 1.**
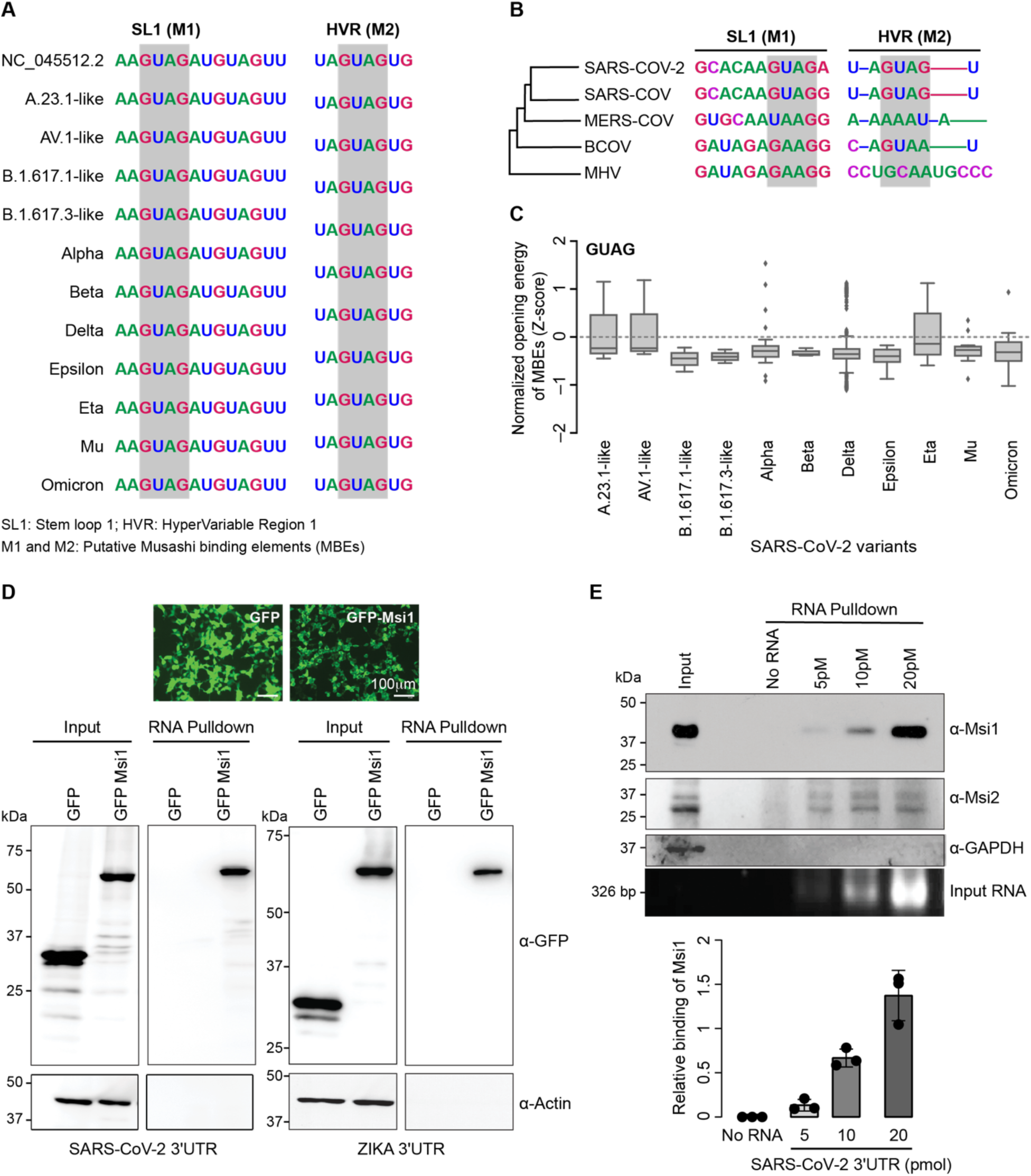
SARS-CoV-2 3’UTR contains conserved Musashi binding elements. **(A)** Multiple sequence alignment of Musashi binding elements (MBE) in coronaviruses. **(B)** MBEs from different variants of SARS-CoV-2 highlighting their conservation status. **(C)** Box plot of distribution of normalised opening energies for (GUAG) Msi consensus site across SARS-CoV-2 variants. **(D)** RNA pull-down assays performed with the 3’ UTRs of SARS-CoV-2 and Zika virus. In vitro transcribed biotinylated 3’ UTR RNA from SARS-CoV-2 and Zika virus were incubated with GFP or GFP-Msi1 expressing HEK293 cell extracts and RNA-protein complexes were captured on streptavidin beads. Representative image of western blots probed with antibodies against Msi1 and Actin as loading control. Panel on top depicts transfection efficiency of GFP and GFP-Msi1. **(E)** RNA pull-down assays were performed with increasing concentrations of 3’ UTR of SARS-CoV-2. Different concentrations of biotinylated RNA were incubated with Caco-2 cell extracts and RNA-protein complexes were captured on streptavidin beads. Representative image of western blots probed with antibodies against Musashi-1 (Msi1), Musashi-2 (Msi2) and Gapdh as control. Corresponding protein and RNA inputs are shown. The graph at the bottom shows a densitometric analysis of Msi1 binding to increasing concentrations of SARS-CoV-2 3’UTR.

### Msi1 binds to 3’UTR of SARS-CoV-2 both *in vitro* and *in vivo*

Since, Msi1 is (i) known to regulate replication of RNA viruses (Zika (3) and HCV (39)) and (ii) involved in intestinal homeostasis (40,41), the organ which serves as a reservoir for SARS-CoV-2, we focussed on the role of Msi1 on the regulation of SARS-CoV-2 infection. To check if Msi1 binds directly to the 3’UTR of SARS-CoV-2, we performed RNA pulldowns with His-tagged Msi1 and the *in vitro* biotinylated SARS-CoV-2 3’UTR. We identified that Msi1 directly binds to SARS-CoV-2 3’UTR at a concentration as low as 5 pmol of RNA and saturated at 10 pmol of RNA with 60 nM of Msi1 protein (**Figure 2A, Supplementary Figure 2A** and 2B). We did not detect any binding of Msi1 with the 5’UTR of SARS-CoV-2 under the same conditions (**Supplementary Figure 2C**). To narrow down the preference of Msi1 binding in the 3’UTR, we next performed Molecular Dynamics (MD) simulations for the two sites (M1 and M2) and compared them to Zika 3’UTR. The variation in the RMSD and RMSF of Msi1 in complex with M2 in the HVR in comparison to Zika 3’UTR was lesser than that of Msi1 in complex with M1 (**Figure 2B, Supplementary Figure 2D**-2E).

**Figure 2.**
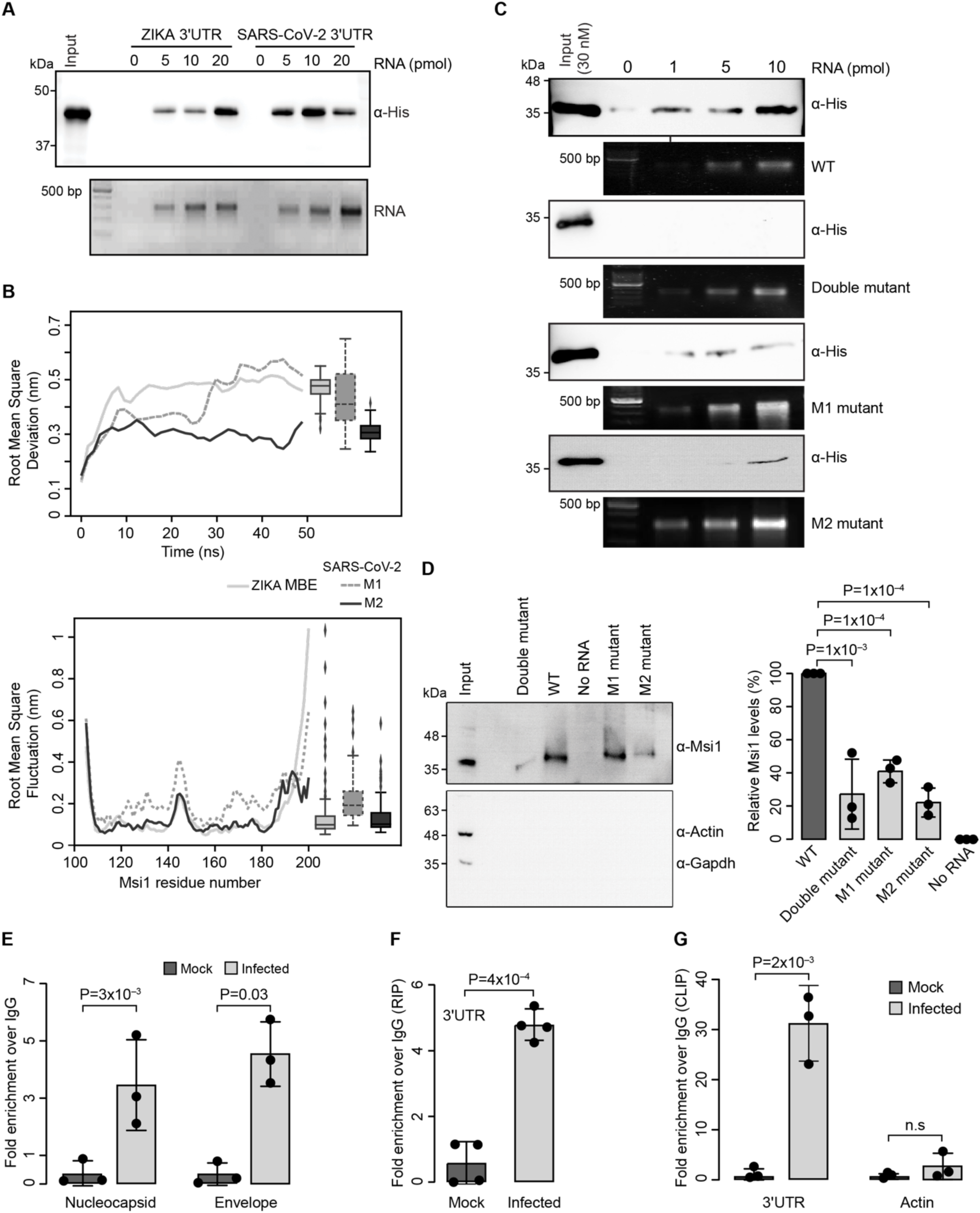
Msi1 directly binds to SARS-CoV-2 3’UTR. **(A)** RNA pull-down of recombinant His-Msi1 protein with different concentrations of in vitro transcribed SARS-CoV-2 3’UTR and Zika virus, followed by immunoblot analysis with His antibody. RNA and protein inputs are depicted. **(B)** Comparison of MD simulations of Zika virus 3’UTR (WT) with the two MBEs in SARS-CoV-2, M1 (SL1) and M2 (HVR1). **(C)** RNA pull down of recombinant His-Msi1 protein with in vitro transcribed wildtype 3’UTR of SARS-CoV-2 and individual mutations in M1, M2 and the double mutant followed by immunoblot analysis probed with His-tag antibody. RNA input for the respective RNA pulldown has been shown below each immunoblot. **(D)** RNA pull-down assays of endogenous Msi1 from Caco-2 lysate with the wildtype 3’UTR, M1, M2 and double mutant of SARS-CoV-2. Representative image of immunoblots probed with antibodies against Msi1 and with Actin and Gapdh as a loading control. Graph on the right provides the densitometric analysis depicting the amount of Msi1 in RNA pull-down for the corresponding 3’UTR of SARS-CoV-2. **(E)** The graphs depict the quantities of Msi1 bound viral Nucleocapsid and Envelope RNAs and **(F)** 3’UTR regions of SARS-CoV-2 genome. upon RNA Immunoprecipitation (RIP) from mock- and SARS-CoV-2 infected Caco-2 cells (Multiplicity of infection; MOI: 1 FFU/cell). Rabbit IgG or Msi1 antibodies were used for immunoprecipitation. RIP values are presented as a percentage of input following subtraction of the IgG background. Nucleocapsid and Envelope transcripts were quantitated by TaqMan assay, whereas 3’UTR was quantified with SYBR qPCR. Bar charts depict mean± SD. n=3,4 biological replicates. P-values were obtained using two-tailed paired Student’s t-test. **(G)** Msi1 CLIP from Caco-2 cells with mock and infected cells as indicated. CLIP was performed with rabbit IgG or Msi1 antibodies. Graphs show qPCRs of bound transcripts. n=3 biological replicates. P-values were obtained using two-tailed paired Student’s t-test; n.s indicates not significant.

To consolidate these findings, we performed site-directed mutagenesis perturbing M1 and M2 individually as well as in combination (**Supplementary Figure 2D**). While disruption of the M2 site in HVR drastically reduced the binding of Msi1 to the 3’UTR, disruption of the M1 site in SL1 showed a modest reduction in binding (**Figure 2C**). The dual mutant (M1 and M2) showed almost complete abrogation of binding indicating that these sites could be the primary contributors to Msi1 binding to SARS-CoV-2 3’UTR. We further validated these findings using endogenous Msi1 protein from Caco-2 cell lysates (**Figure 2D**). To examine if the Msi1 binding to SARS-CoV-2 3’UTR occurs *in vivo* during viral infection, we performed RNA immunoprecipitations in Caco-2 cell lines infected with SARS-CoV-2. Using viral Nucleocapsid and Envelope transcripts and the viral 3’UTR as probes, we showed that upon infection, Msi1 immunoprecipitated SARS-CoV-2 genomic RNA (**Figures 2E-2G**). Crosslinking immunoprecipitation (CLIP) established the direct interaction of Msi1 with the 3’UTR of the SARS-CoV-2 genome (**Figure 2G**). Taken together, these findings establish the binding of Msi1 to SARS-CoV-2 3’UTR, at the two Musashi binding elements present in the SL1 and HVR.

### Depletion of Msi1 levels increases viral load

To establish the physiological relevance of Msi1 interaction with SARS-CoV-2 3’UTR, we generated Msi1 knockouts (KO) in Caco-2 cells using lentivirus-based CRISPR-Cas9. Importantly, the protein levels of Msi1 remain unchanged upon different time points of SARS-CoV-2 infection in Caco-2 cells **(Supplementary Figure 3A)**. Subsequently, we generated the Msi1 KOs and verified the protein levels of both Musashi proteins, Msi1 and Msi2 in the KO pools as well as in individual knock-outs. The guides specifically targeted Msi1 and we could not detect any change in Msi2 protein levels upon loss of Msi1 (**Figure 3A**). We then subjected the parental cells expressing Cas9 and Msi1 knockout pool and the individual clones to SARS-CoV-2 infection (Multiplicity of infection, MOI=1). Viral transcript analysis post-infection revealed a significant increase in the transcript levels of Envelope and RdRp across the Msi1 KO pool and the individual clones, suggesting an increase in viral replication (**Figure 3B**). This increase in viral replication was validated by increased dsRNA positivity in the Msi1 KO cells compared to the control cells (**Figure 3C**). We then performed western blot analysis and compared the nucleocapsid levels between control and knockout cells at 24h and 48h post-infection. We observed a ∼2-3-fold increase in nucleocapsid protein levels in the knock-outs starting from 24h time point, peaking at 48h time point to ∼4 folds (**Figure 3D**). Ileum and colon are known to show elevated expression of the entry receptors for viruses, specifically ACE2, alluding to enhanced infection of the intestinal cells (42,43). To examine the effect of Msi1 on ACE2 levels, we performed immunoblot assay and found that there was no change in ACE2 expression in Msi1 KOs upon infection (**Figure 3D**). This suggests that Msi1 might not regulate the entry of viruses into cells by modulating ACE2 levels. To confirm the infectivity of the virus, we performed a plaque formation assay to check the differences in infectious viral titres between control and knock-out cells. We observed a significant increase in plaque formation in the Msi1 knockouts compared to the control, indicating an increase in infective viral load (**Figure 3E**). While these findings of Msi1 repressing viral load in Caco-2 cells could be attributed to high levels of expression of Msi1 in Caco-2 cells, we wanted to examine the effect of Msi1 in a more physiological setting.

**Figure 3.**
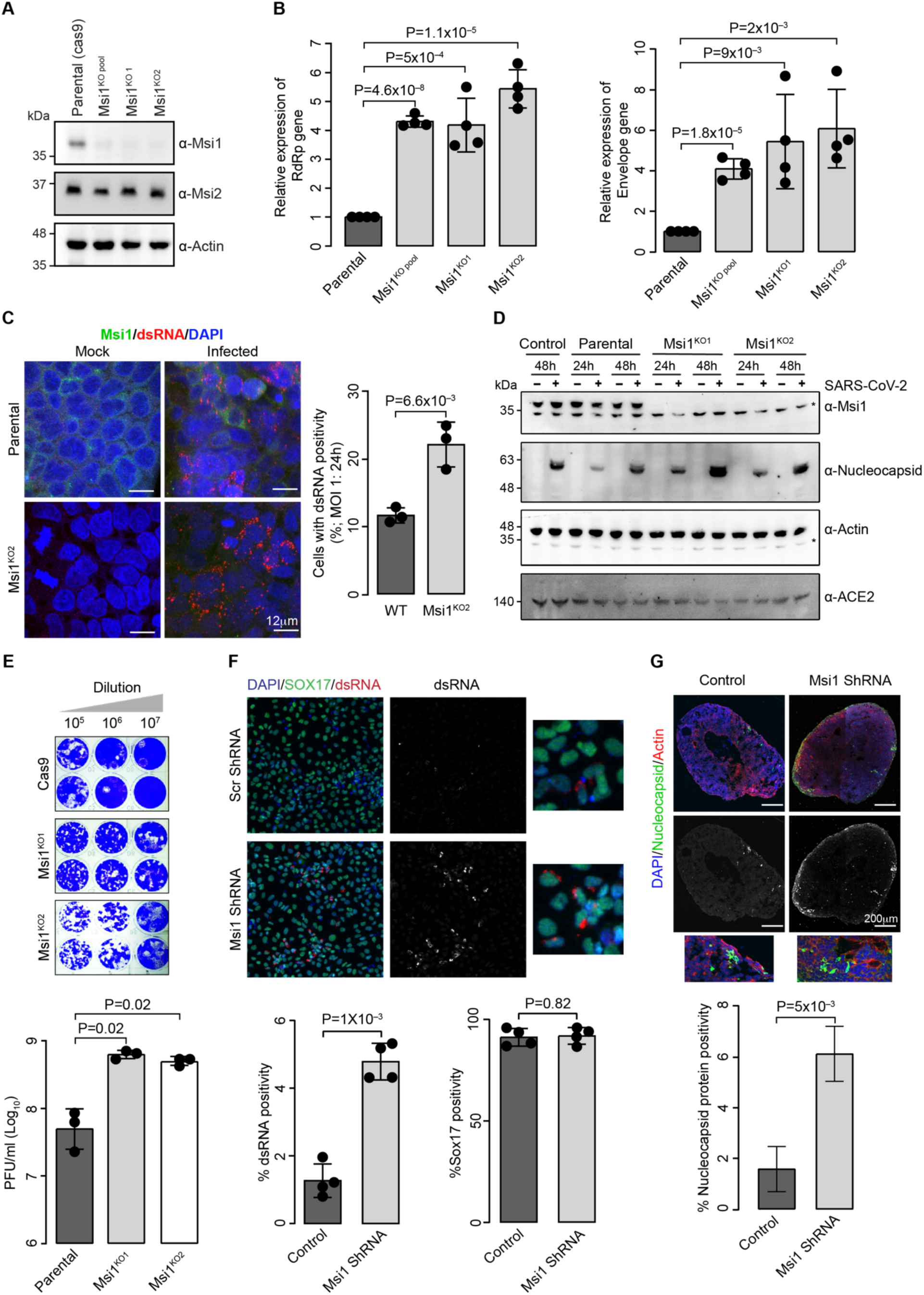
Msi1 depletion promotes SARS-CoV-2 infection. **(A)** Western blot analysis of Caco-2 knockouts (KO) depicting the absence of Msi1 protein and unchanged Msi2 levels. Actin was used as a loading control. **(B)** The graph shows viral RNA copies in Msi1 KO pool or individual KO clones 1 and 2 following infection with SARS CoV2 (MOI: 1 FFU/cell, 48h). In all viral replication assays, RdRp and Envelope transcripts were quantified using TaqMan assay as described in Materials and Methods. N=4 biological replicates. P-values were estimated using two-tailed paired Student’s t-test. **(C)** Confocal microscopy images of SARS CoV2-infected parental and KO2 Caco-2 cells immunostained with antibodies against dsRNA (red), Msi1(green) and DAPI (blue) following mock or SARS CoV2 (MOI: 1 FFU/cell, 48h). **(D)** Western blots of Control, Parental, Msi1 KO cell lines with indicated genotypes, mock- or SARS-CoV-2 infected (MOI:1) across the indicated time points. Blots were probed with antibodies as indicated. Actin was used as a loading control. * indicates non-specific band **(E)** Panel in top shows representative image of Plaques. Graph at the bottom shows relative fold change in the infectious viral titres of SARS-CoV-2 in Parental or Msi1 KO samples compared to those treated with vehicle, represented as fold change in PFU/mL. **(F)** Confocal images of definitive endoderm cells treated with Scrambled or Msi1 shRNA subjected to SARS-CoV-2 infection (MOI:1, 48h). SOX17 depicts DE cells, dsRNA represents viral replication and DAPI the nuclei. The graphs depict quantification of dsRNA and SOX17 positivity in each condition. **(G)** Confocal microscopy images of SARS CoV2-infected Scrambled shRNA or Msi1 shRNA infected gut organoids immunostained with antibodies against Nucleocapsid protein (green), Actin (red) and DAPI (blue). Graph depicts quantification of Nucleocapsid protein positivity normalised to DAPI.

Previous studies have demonstrated that intestinal organoids are susceptible to SARS-CoV-2 infection, with enterocytes being the primary targets, resulting in the production of infectious viral particles and damage of intestinal epithelium (7,44). The Musashi (Msi) family of proteins plays a critical role in maintaining intestinal regeneration and homeostasis after insults by driving the exit of intestinal stem cells from quiescence to enter cell cycle (45). In this context, here, we first assessed the impact of Msi1 depletion on SARS-CoV-2 infection using definitive endoderm (DE) cells, the precursors of intestinal cell lineages. To achieve this, we utilized lentiviral vectors encoding either Msi1-specific shRNA or scrambled shRNA to knock down Msi1 expression in DE cells, followed by infection with SARS-CoV-2. Immunofluorescence staining for double-stranded RNA (dsRNA), a marker of viral replication, and Sox17, a DE marker, were performed. While only 1% of DE cells were positive for dsRNA at a multiplicity of infection (MOI) of 1, approximately 5% of Msi1-depleted DE cells exhibited dsRNA positivity. Importantly, Sox17 expression was largely unaffected by Msi1 depletion, indicating that the effect of Msi1 on stem cell population remains unchanged while only the infection of cells was affected. These findings led us to further investigate the role of Msi1 in regulating SARS-CoV-2 infection within intestinal organoids. For this, we generated mid-hindgut organoids from induced pluripotent stem cells (iPSCs) and depleted Msi1 using shRNA (iPSCs; **Figure 3F, Supplementary Figure 3B**). Notably, we observed a significant increase in nucleocapsid and dsRNA positivity in Msi1-depleted organoids compared to control organoids (**Figure 3G, Supplementary Figure 3C**-3D). Collectively, these results suggest that Msi1 acts as a negative regulator of SARS-CoV-2 infection both *in vitro* and *in vivo*, highlighting its potential role in modulating viral susceptibility in the intestinal epithelium.

Given that Msi1 knockout (KO) cells exhibited a significant increase in viral titres, we reintroduced the Msi1 transgene into the knockout background to assess whether the inhibitory effect of Msi1 could be restored (**Figure 4A)**. Upon re-expression of Msi1 in the KO background, we observed a 50% reduction in the expression levels of the viral Envelope and RdRp transcripts (**Figure 4B)**. This result was corroborated by a plaque assay, which showed a reduction in cytopathic effects (CPEs) upon Msi1 reintroduction (**Figure 4C)**. Additionally, immunoblot analysis of cells following transgene expression and subsequent infection demonstrated decreased levels of viral nucleocapsid protein in the presence of Msi1, confirming the inhibitory role of Msi1 in suppressing viral replication (**Figure 4D**).

**Figure 4.**
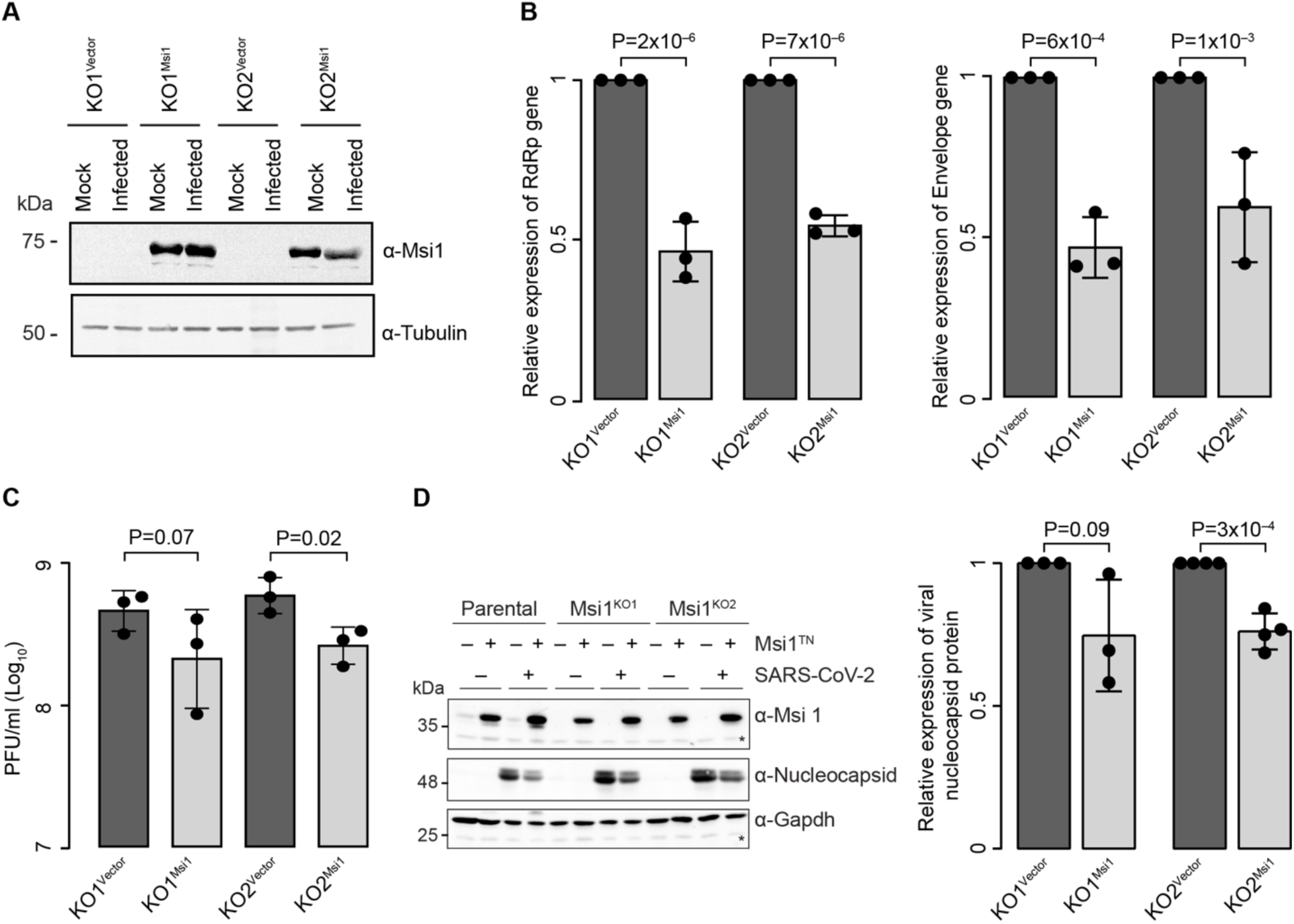
Msi1 transgene attenuates the replication of SARS-CoV-2 in cells. **(A)** Western blot depicting the expression of GFP (KO^Vector^) or Msi1-GFP (KO^Msi1^) transgene in the Msi1 KO background. Tubulin was used as loading control. **(B)** Distribution of viral RNA copies in Msi1 KO cells rescued with empty vector or Msi1 transgene following infection with SARS-CoV-2 (MOI: 1 FFU/cell, 48h). n=3 biological replicates. P-values were estimated using two-tailed paired Student’s t-test. **(C)** Relative fold change in the infectious viral titres of SARS-CoV-2 in Msi1 KO cells rescued with Vector or Msi1. **(D)** Western blots of cell lines with indicated genotypes mock-infected or infected with SARS-CoV-2 (MOI:1, 48h) for the indicated time points (left panel). Blots were probed with antibodies as indicated, with Gapdh serving as a loading control. Graph on the right depicts quantification of the nucleocapsid protein in the transgene rescued Msi1 cell lines.

### Msi1 mediates SARs-CoV-2 inhibitory effects through repression of viral translation

Given that Msi1 possesses two conserved RRMs capable of interacting with RNA, we next examined whether these minimal RNA recognition motifs are sufficient to recapitulate the effects of the full-length protein on SARS-CoV-2. We performed a rescue experiment in Caco-2 cells expressing RRMs (Msi1 20-190) along with GFP and GFP-Msi1 and examined the changes in the levels of viral transcripts **(Figure 5A)**. Surprisingly, the RRMs alone did not show any significant change in the transcript levels of the viral genes, indicating that the activity of the intact Msi1 protein is required for the observed inhibitory effects on the virus **(Figure 5B)**. Since Msi1 is a known translational repressor, we investigated if its binding to viral 3’ UTR could impact the viral translation. For this, we performed a luciferase reporter assay by transfecting the 3’UTR of SARS-CoV-2 along with Msi1. Overexpression of Msi1 led to a 20%-30% reduction in the expression of the firefly luciferase/Renilla luciferase reporter from the wild-type 3’UTR. However, we did not observe any significant difference in reporter expression in the MBE double mutant 3’UTR (**Figure 5C**). Importantly, the Msi1 gene construct with only the two RNA recognition motifs was not sufficient for repressing the reporter driven translation (**Figure 5D)**. These findings suggest that full length Msi1 is essential to repress translation of viral transcripts.

**Figure 5.**
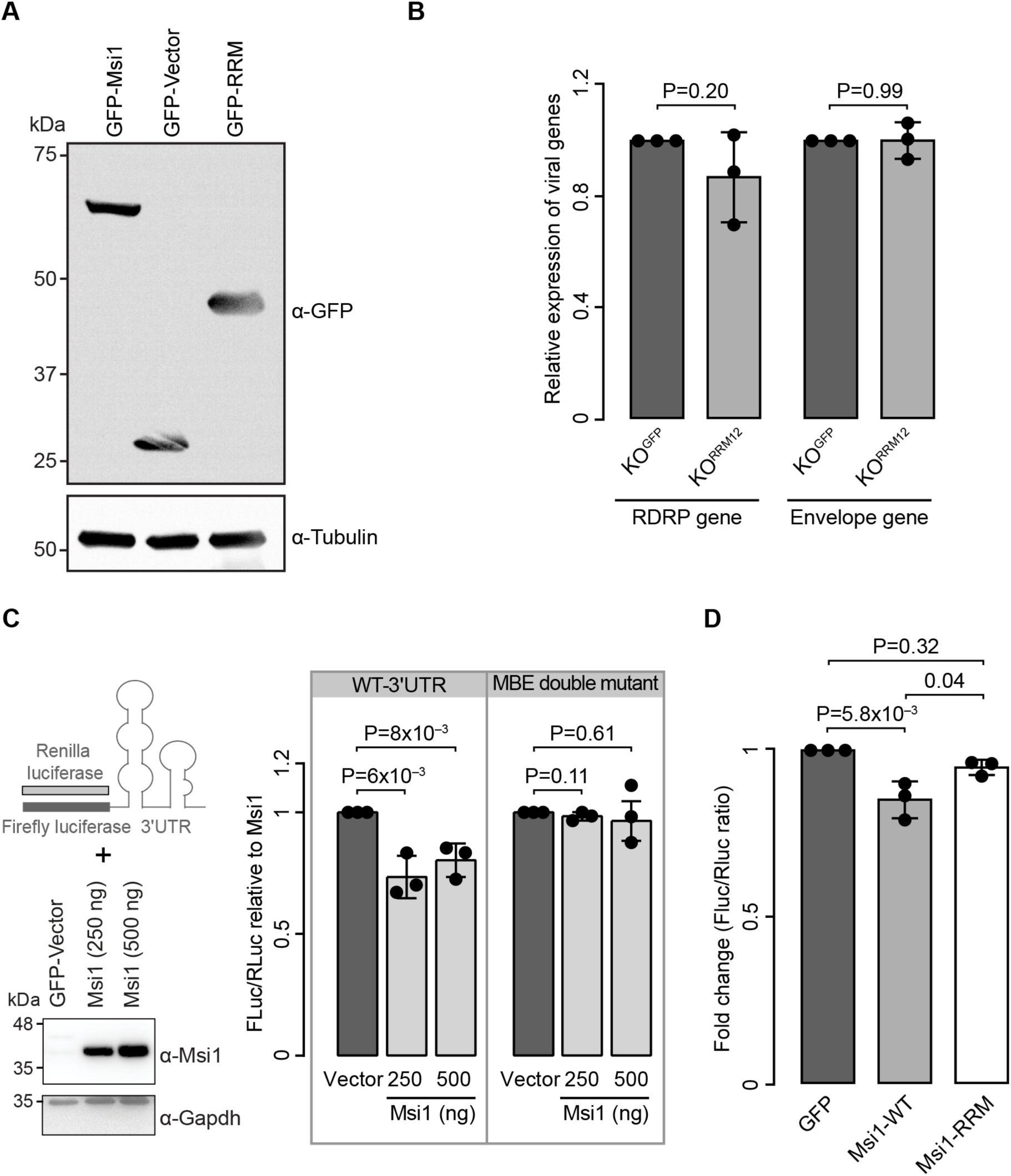
Msi1 represses viral translation by binding to the 3’UTR of SARS-COV-2. **(A)** Western blot depicting expression of GFP, Msi1-GFP, RRM-GFP (Msi1 1-190), along with tubulin as loading control. **(B)** Graph showing relative expression of RdRp and Envelope gene of SARS-CoV-2 in GFP or GFP-RRM expressed Msi1 KO cells. **(C)** Schematic representation of luciferase assay using 3’UTR of SARS-CoV-2. Western blot depicts the dose-dependent expression of Msi1 in HEK293T cells along with Gapdh as loading control. Graph on the right shows the fold change in firefly luciferase/Renilla luciferase reporter. **(D)** Graph depicts fold change Firefly /Renilla luciferase ratio after transfection with GFP, Msi1 GFP or GFP RRM. P-values were estimated using two-tailed paired Student’s t-test.

### Msi1 represses PABP1-mediated translation of SARS-CoV-2 minigene

Given that the interactions between the 5’ UTR and 3’ UTRs, along with their associated molecular interactions, are known to influence viral replication and translation, we performed a SARS-CoV-2 minigene reporter assay. In this construct, a Gaussia luciferase reporter is flanked by 5’ and 3’UTR of SARS-CoV-2 and a segment of Nsp1. This construct closely mimics the viral replicon, which can be used as a surrogate for viral translation (18). Once we confirmed that the expression levels of the wild type and the double mutant minigene transcripts were not significantly different, we performed translation analysis using Gaussia reporter as surrogate (**Figure 6A**). While we observed a 30-40% Msi1-driven decrease in Gaussia luciferase expression in a dose-dependent manner (**Figure 6B**), the double mutant 3’UTR, was not responsive to the Msi1 regulated Gaussia luciferase expression. Taken together, these observations suggest that Msi1 binding to the M1 and M2 sites of SARS-CoV-2 3’UTR represses translation. This suggests that Msi1 may reduce viral load through 3’ UTR-driven translational repression in vivo. Notably, SARS-CoV-2 RNAs are polyadenylated, which can influence their stability and translation (46). For several coronaviruses, the interaction between the poly (A) tail and poly(A)-binding protein 1 (PABP1) has been shown to regulate viral translation (47,48). It is well established that Msi1 competes with PABP1, leading to translational repression (49). To investigate whether Msi1 mediates SARS-CoV-2 translational repression through this mechanism, we co-transfected PABP1 with the minigene and observed a 1.5- to 2.5-fold increase in translation (**Figure 6C**). However, when Msi1 was co-transfected, this upregulation was reversed, indicating that Msi1 can inhibit PABP1-mediated translation of SARS-CoV-2 (**Figure 6D)**.

**Figure 6.**
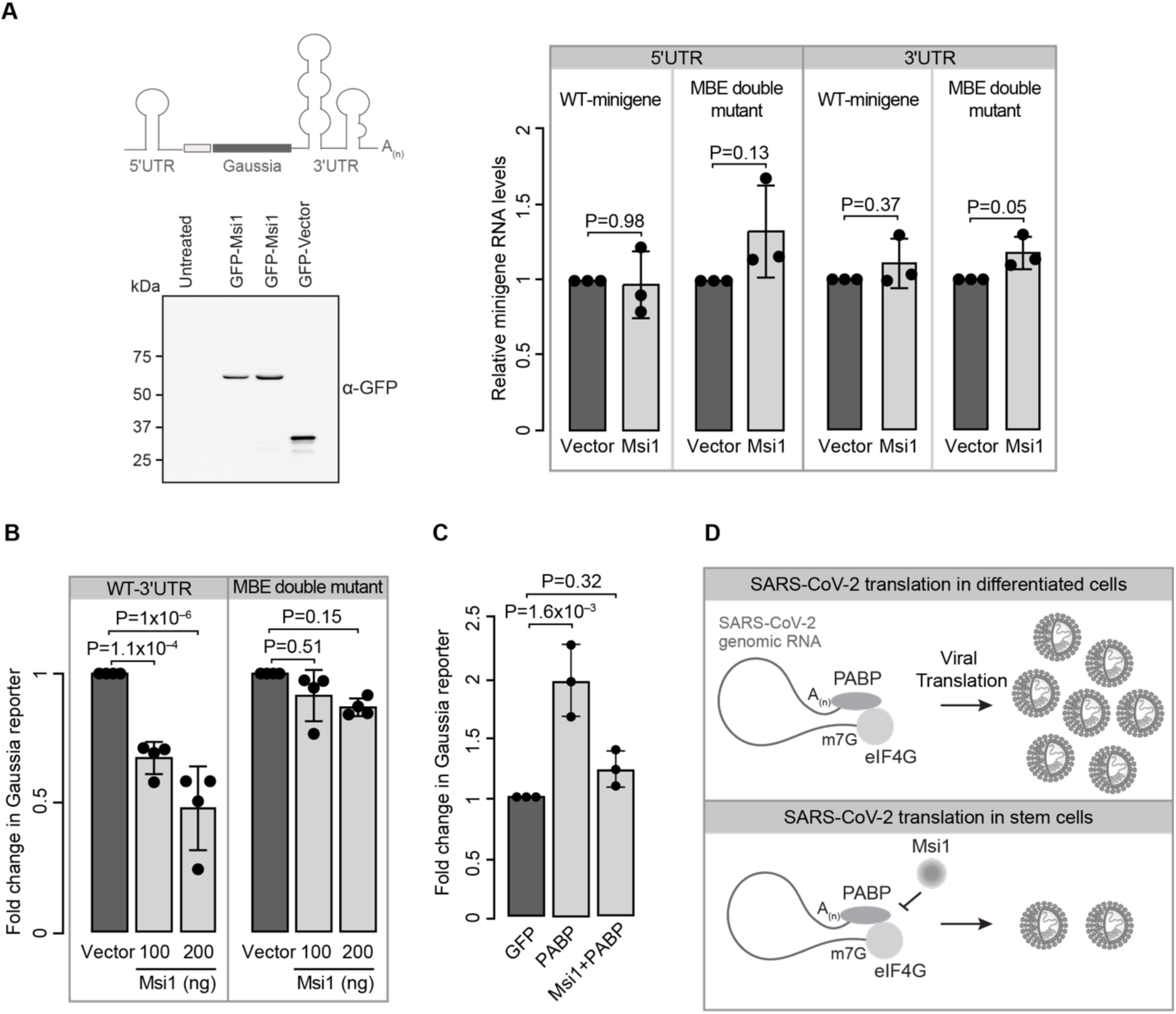
Msi1 represses PABP mediated viral translation. **(A)** Schematic representation of the minigene assay with the Gaussia reporter cloned between 3’UTR and 5’UTR of SARS-COV-2. Western blot below depicts the dose dependent expression of GFP-Msi1 (Msi1) and GFP transgene (vector). Graph on the right depicts relative expression of wildtype and MBE-double mutant minigene transcript transfected for Gaussia luciferase assay in presence of GFP (Vector) and GFP-Msi1 (Msi1). (**B**) The graph represents fold change in Gaussia reporter upon transfection of the minigene containing wildtype 3’UTR or the MBE double mutant indicated plasmids. P-values were estimated using two-tailed paired Student’s t-test. **(C)** Quantification of change in WT minigene Gaussia reporter level upon transfection of GFP, PABP and Msi1+PABP. P-values were estimated using two-tailed paired Student’s t-test. (**D)** Schematic representation of Msi1 binding to the 3’UTR and inhibiting viral translation in stem cells by possibly interfering with the PABP binding and translation of the viral proteins resulting in lesser number viral particles.

## DISCUSSION

Previous studies have shown that Msi1 promotes replication of Zika virus, albeit the underlying molecular mechanism is not yet known (3,50). Here, we show the translational repressor function of Msi1 in a viral context. Similar to other RNA viruses, SARS-CoV-2 engages both canonical and non-canonical pathways of translation (51). The 3’UTR of coronaviruses contains a number of cis-acting elements which are essential for RNA synthesis and translation. Moreover, the hypervariable region (HVR) which contains stem loop motif II (S2M) is a binding hub for host translation factors (18). In this context, since Msi1 binds in the close vicinity to HVR, it is likely that Msi1 can compete for binding with other translational regulators and inhibit translation of SARS-CoV-2. For instance, here we show that the Msi1 can repress SARs-CoV-2 translation by inhibiting PABP (49). The minigene assay encompassing 5’UTR and 3’UTR with Gaussia reporter mimics the viral genomic RNA and shows that Msi1 binds to 3’UTR and represses translation of SARS-CoV-2 genomic RNA in a dose-dependent manner. While the mere use of 3’UTR in the reporter assay shows a 25-30% downregulation, usage of complete viral minigene shows a 40-60% reduction, possibly because the minigene constructs better mimics the *in vivo* circularization of the genome.

Recent studies highlight that targeting the 3’UTR, which is conserved across most variants of SARS-CoV-2, could be used as an intervention strategy (24,52). Extensive RNA-protein interaction characterization of SARs-CoV-2 across various human cell lines has been done recently (22,53,54). While Msi1 has been identified as a potential interactor in Huh7 cells, Msi1 has not been detected as an interactor in other cell lines, such as HEK293 and Calu3. The potential reasons for this discrepancy could be (i) cells do not express endogenous Msi1, as is the case in HEK293 (3), (ii) downregulation of Msi1 protein upon SARs-CoV-2 infection, as shown here in both Calu3 and Huh7 cell lines and (iii) expression of truncated forms of Msi1, for instance, the Msi1 expressed in Calu3 lacks the C-terminus, which might influence the viral RNA-host protein interaction (**Supplementary Figure 5A**). Furthermore, unlike Zika, wherein only Msi1 is bound to the 3’UTR, both Msi1 and Msi2 are shown to bind to SARS-CoV-2 3’UTR. This is noteworthy since, Msi1 and Msi2 are thought to be functionally redundant, although their expression pattern varies. For instance, while Msi1 is thought to be a stem cell marker, Msi2 is found to be more ubiquitous (41). Indeed a single-cell gene expression analysis of intestinal cells revealed that Msi1 expression is restricted to Transient Amplifying (TA) cells and stem cells, while Msi2 has a widespread expression across cell types, with the highest being in the cycling TA population. Moreover, enterocytes that are readily infected by SARS-CoV-2 show negligible expression of Msi1 (**Supplementary Figure 5B**). Likewise staining of cerebral organoids showed neural progenitors express high levels of Msi1 compared to Msi2 (**Supplementary Figure 5C**). This is interesting since neural progenitors have been shown to be less amenable to SARS-CoV-2 infection (55).

To recapitulate physiology better, we performed an analysis of single-cell RNAseq data of intestinal organoids infected with SARS-CoV-2 and found that at a 12h time point, Msi1 transcript significantly goes down (**Supplementary Figure 5D**). Integrating our findings pertaining to Msi1 mediated repression of SARs-CoV-2 translation and that of single-cell RNA sequencing data of intestinal organoids upon viral infection, it is tempting to speculate that the virus most likely downregulates Msi1 during the onset of its gRNA translation. However, when the innate immune response sets in at 24h time point, enhanced expression of Msi1 is observed. This is interesting in the context of the expression pattern of Msi1, which is primarily expressed in stem cells. It is now well known that different stem cells have cell type-specific groups of interferon-stimulated genes, which makes the stem cells resistant to viral infection compared to their differentiated progeny (56–58). For instance, in addition to directly binding and repressing translation, Msi1 could also activate specific modules like Protein Kinase R, which are activated by dsRNA and can restrict viral translation by inducing a shutdown (59–61). Thus, an interplay between innate immune response and RBPs such as Msi1 could play vital roles in viral containment, which needs to be further explored.

## DATA AVAILABILITY

All data are available in the main text or the supplementary materials.

## AUTHOR CONTRIBUTIONS

Conceptualization, P.L.C., S.G., S.B., S.C; Methodology, P.L.C., S.G., R.V.K., S.B, S.C., A.G.K., K.H.H., D.G., D.T.D., and R.R.; Formal analysis, S.G., D.G., D.T., R.R and P.L.C.; Visualization, P.L.C, S.G, R.V.K, S.C. and D.G.; Writing– original draft, P.L.C., S.G and S.C.; Writing– review & editing, P.L.C., S.G., K.H.H., D.G., D.T., R.V.K., and S.C.; Supervision, P.L.C, and K.H.H.

## ACKNOWLEDGEMENTS

We thank Jingxin Wang Lab for the Gaussia minigene construct and Amit Kumar for assistance with the logistics in the BSL3 lab. Stem cell studies were performed after obtaining due approval from the Institute committee for stem cell research (IC-SCR), CCMB (ICSCR-51/2022-R1/2023). SARS-CoV-2 isolation and culturing was approved by the Institutional Biosafety Committee of CCMB.

## REFERENCES

1. Zhang, F., Chase-Topping, M., Guo, C.G., van Bunnik, B.A.D., Brierley, L. and Woolhouse, M.E.J. (2020) Global discovery of human-infective RNA viruses: A modelling analysis. PLoS Pathog, 16, e1009079.

2. Lisy, S., Rothamel, K. and Ascano, M. (2021) RNA Binding Proteins as Pioneer Determinants of Infection: Protective, Proviral, or Both? Viruses, 13.

3. Chavali, P.L., Stojic, L., Meredith, L.W., Joseph, N., Nahorski, M.S., Sanford, T.J., Sweeney, T.R., Krishna, B.A., Hosmillo, M., Firth, A.E. et al. (2017) Neurodevelopmental protein Musashi-1 interacts with the Zika genome and promotes viral replication. Science, 357, 83–88.

4. Darai, N., Mahalapbutr, P., Wolschann, P., Lee, V.S., Wolfinger, M.T. and Rungrotmongkol, T. (2022) Theoretical studies on RNA recognition by Musashi 1 RNA-binding protein. Sci Rep, 12, 12137.

5. Schneider, A.B. and Wolfinger, M.T. (2019) Musashi binding elements in Zika and related Flavivirus 3’UTRs: A comparative study in silico. Sci Rep, 9, 6911.

6. Chen, X., Wang, Y., Xu, Z., Cheng, M.L., Ma, Q.Q., Li, R.T., Wang, Z.J., Zhao, H., Zuo, X., Li, X.F. et al. (2023) Zika virus RNA structure controls its unique neurotropism by bipartite binding to Musashi-1. Nat Commun, 14, 1134.

7. Lamers, M.M., Beumer, J., van der Vaart, J., Knoops, K., Puschhof, J., Breugem, T.I., Ravelli, R.B.G., Paul van Schayck, J., Mykytyn, A.Z., Duimel, H.Q., et al. (2020) SARS-CoV-2 productively infects human gut enterocytes. Science, 369, 50–54.

8. Tang, H., Hammack, C., Ogden, S.C., Wen, Z., Qian, X., Li, Y., Yao, B., Shin, J., Zhang, F., Lee, E.M. et al. (2016) Zika Virus Infects Human Cortical Neural Progenitors and Attenuates Their Growth. Cell Stem Cell, 18, 587–590.

9. Zhang, B.Z., Chu, H., Han, S., Shuai, H., Deng, J., Hu, Y.F., Gong, H.R., Lee, A.C., Zou, Z., Yau, T. et al. (2020) SARS-CoV-2 infects human neural progenitor cells and brain organoids. Cell Res, 30, 928–931.

10. Lan, J., Ge, J., Yu, J., Shan, S., Zhou, H., Fan, S., Zhang, Q., Shi, X., Wang, Q., Zhang, L. et al. (2020) Structure of the SARS-CoV-2 spike receptor-binding domain bound to the ACE2 receptor. Nature, 581, 215–220.

11. Chen, T.H., Hsu, M.T., Lee, M.Y. and Chou, C.K. (2022) Gastrointestinal Involvement in SARS-CoV-2 Infection. Viruses, 14.

12. Cheung, K.S., Hung, I.F.N., Chan, P.P.Y., Lung, K.C., Tso, E., Liu, R., Ng, Y.Y., Chu, M.Y., Chung, T.W.H., Tam, A.R. et al. (2020) Gastrointestinal Manifestations of SARS-CoV-2 Infection and Virus Load in Fecal Samples From a Hong Kong Cohort: Systematic Review and Meta-analysis. Gastroenterology, 159, 81–95.

13. Mao, L., Jin, H., Wang, M., Hu, Y., Chen, S., He, Q., Chang, J., Hong, C., Zhou, Y., Wang, D. et al. (2020) Neurologic Manifestations of Hospitalized Patients With Coronavirus Disease 2019 in Wuhan, China. JAMA Neurol, 77, 683–690.

14. Sharma, O., Sultan, A.A., Ding, H. and Triggle, C.R. (2020) A Review of the Progress and Challenges of Developing a Vaccine for COVID-19. Front Immunol, 11, 585354.

15. Hoffmann, M., Kleine-Weber, H., Schroeder, S., Kruger, N., Herrler, T., Erichsen, S., Schiergens, T.S., Herrler, G., Wu, N.H., Nitsche, A. et al. (2020) SARS-CoV-2 Cell Entry Depends on ACE2 and TMPRSS2 and Is Blocked by a Clinically Proven Protease Inhibitor. Cell, 181, 271–280 e278.

16. Jackson, C.B., Farzan, M., Chen, B. and Choe, H. (2022) Mechanisms of SARS-CoV-2 entry into cells. Nat Rev Mol Cell Biol, 23, 3–20.

17. Sola, I., Almazan, F., Zuniga, S. and Enjuanes, L. (2015) Continuous and Discontinuous RNA Synthesis in Coronaviruses. Annu Rev Virol, 2, 265–288.

18. Zhao, J., Qiu, J., Aryal, S., Hackett, J.L. and Wang, J. (2020) The RNA Architecture of the SARS-CoV-2 3’-Untranslated Region. Viruses, 12.

19. Flynn, R.A., Belk, J.A., Qi, Y., Yasumoto, Y., Wei, J., Alfajaro, M.M., Shi, Q., Mumbach, M.R., Limaye, A., DeWeirdt, P.C. et al. (2021) Discovery and functional interrogation of SARS-CoV-2 RNA-host protein interactions. Cell, 184, 2394–2411 e2316.

20. Kamel, W., Noerenberg, M., Cerikan, B., Chen, H., Jarvelin, A.I., Kammoun, M., Lee, J.Y., Shuai, N., Garcia-Moreno, M., Andrejeva, A. et al. (2021) Global analysis of protein-RNA interactions in SARS-CoV-2-infected cells reveals key regulators of infection. Mol Cell, 81, 2851–2867 e2857.

21. Labeau, A., Fery-Simonian, L., Lefevre-Utile, A., Pourcelot, M., Bonnet-Madin, L., Soumelis, V., Lotteau, V., Vidalain, P.O., Amara, A. and Meertens, L. (2022) Characterization and functional interrogation of the SARS-CoV-2 RNA interactome. Cell Rep, 39, 110744.

22. Lee, S., Lee, Y.S., Choi, Y., Son, A., Park, Y., Lee, K.M., Kim, J., Kim, J.S. and Kim, V.N. (2021) The SARS-CoV-2 RNA interactome. Mol Cell, 81, 2838–2850 e2836.

23. Jiang, L., Xiao, M., Liao, Q.Q., Zheng, L., Li, C., Liu, Y., Yang, B., Ren, A., Jiang, C. and Feng, X.H. (2023) High-sensitivity profiling of SARS-CoV-2 noncoding region-host protein interactome reveals the potential regulatory role of negative-sense viral RNA. mSystems, 8, e0013523.

24. Rangan, R., Zheludev, I.N., Hagey, R.J., Pham, E.A., Wayment-Steele, H.K., Glenn, J.S. and Das, R. (2020) RNA genome conservation and secondary structure in SARS-CoV-2 and SARS-related viruses: a first look. RNA, 26, 937–959.

25. Lorenz, R., Bernhart, S.H., Honer Zu Siederdissen, C., Tafer, H., Flamm, C., Stadler, P.F. and Hofacker, I.L. (2011) ViennaRNA Package 2.0. Algorithms Mol Biol, 6, 26.

26. Pronk, S., Pall, S., Schulz, R., Larsson, P., Bjelkmar, P., Apostolov, R., Shirts, M.R., Smith, J.C., Kasson, P.M., van der Spoel, D., et al. (2013) GROMACS 4.5: a high-throughput and highly parallel open source molecular simulation toolkit. Bioinformatics, 29, 845–854.

27. Van Der Spoel, D., Lindahl, E., Hess, B., Groenhof, G., Mark, A.E. and Berendsen, H.J. (2005) GROMACS: fast, flexible, and free. J Comput Chem, 26, 1701–1718.

28. Xu, Y., Vanommeslaeghe, K., Aleksandrov, A., MacKerell, A.D., Jr. and Nilsson, L. (2016) Additive CHARMM force field for naturally occurring modified ribonucleotides. J Comput Chem, 37, 896–912.

29. York, D.M., Darden, T.A., Pedersen, L.G. and Anderson, M.W. (1993) Molecular dynamics simulation of HIV-1 protease in a crystalline environment and in solution. Biochemistry, 32, 1443–1453.

30. Hess, B. (2008) P-LINCS: A Parallel Linear Constraint Solver for Molecular Simulation. J Chem Theory Comput, 4, 116–122.

31. van Gunsteren, W.F., Berendsen, H.J., Hermans, J., Hol, W.G. and Postma, J.P. (1983) Computer simulation of the dynamics of hydrated protein crystals and its comparison with x-ray data. Proc Natl Acad Sci U S A, 80, 4315–4319.

32. Bellaousov, S., Reuter, J.S., Seetin, M.G. and Mathews, D.H. (2013) RNAstructure: Web servers for RNA secondary structure prediction and analysis. Nucleic Acids Res, 41, W471–474.

33. Mathews, D.H., Sabina, J., Zuker, M. and Turner, D.H. (1999) Expanded sequence dependence of thermodynamic parameters improves prediction of RNA secondary structure. J Mol Biol, 288, 911–940.

34. Mathews, D.H., Disney, M.D., Childs, J.L., Schroeder, S.J., Zuker, M. and Turner, D.H. (2004) Incorporating chemical modification constraints into a dynamic programming algorithm for prediction of RNA secondary structure. Proc Natl Acad Sci U S A, 101, 7287–7292.

35. Johnson, P.Z., Kasprzak, W.K., Shapiro, B.A. and Simon, A.E. (2019) RNA2Drawer: geometrically strict drawing of nucleic acid structures with graphical structure editing and highlighting of complementary subsequences. RNA Biol, 16, 1667–1671.

36. Sarkar, G. and Sommer, S.S. (1990) The “megaprimer” method of site-directed mutagenesis. Biotechniques, 8, 404–407.

37. Gupta, D., Parthasarathy, H., Sah, V., Tandel, D., Vedagiri, D., Reddy, S. and Harshan, K.H. (2021) Inactivation of SARS-CoV-2 by beta-propiolactone causes aggregation of viral particles and loss of antigenic potential. Virus Res, 305, 198555.

38. Goebel, S.J., Miller, T.B., Bennett, C.J., Bernard, K.A. and Masters, P.S. (2007) A hypervariable region within the 3’ cis-acting element of the murine coronavirus genome is nonessential for RNA synthesis but affects pathogenesis. J Virol, 81, 1274–1287.

39. Rios-Marco, P., Romero-Lopez, C. and Berzal-Herranz, A. (2016) The cis-acting replication element of the Hepatitis C virus genome recruits host factors that influence viral replication and translation. Sci Rep, 6, 25729.

40. Lan, S.Y., Yu, T., Xia, Z.S., Yuan, Y.H., Shi, L., Lin, Y., Huang, K.H. and Chen, Q.K. (2010) Musashi 1-positive cells derived from mouse embryonic stem cells can differentiate into neural and intestinal epithelial-like cells in vivo. Cell Biol Int, 34, 1171–1180.

41. Li, N., Yousefi, M., Nakauka-Ddamba, A., Li, F., Vandivier, L., Parada, K., Woo, D.H., Wang, S., Naqvi, A.S., Rao, S. et al. (2015) The Msi Family of RNA-Binding Proteins Function Redundantly as Intestinal Oncoproteins. Cell Rep, 13, 2440–2455.

42. Guo, Y., Wang, B., Gao, H., Gao, L., Hua, R. and Xu, J.D. (2021) ACE2 in the Gut: The Center of the 2019-nCoV Infected Pathology. Front Mol Biosci, 8, 708336.

43. Zhang, H., Kang, Z., Gong, H., Xu, D., Wang, J., Li, Z., Li, Z., Cui, X., Xiao, J., Zhan, J. et al. (2020) Digestive system is a potential route of COVID-19: an analysis of single-cell coexpression pattern of key proteins in viral entry process. Gut, 69, 1010–1018.

44. Han, Y., Yang, L., Lacko, L.A. and Chen, S. (2022) Human organoid models to study SARS-CoV-2 infection. Nat Methods, 19, 418–428.

45. Yousefi, M., Li, N., Nakauka-Ddamba, A., Wang, S., Davidow, K., Schoenberger, J., Yu, Z., Jensen, S.T., Kharas, M.G. and Lengner, C.J. (2016) Msi RNA-binding proteins control reserve intestinal stem cell quiescence. J Cell Biol, 215, 401–413.

46. Kim, D., Lee, J.Y., Yang, J.S., Kim, J.W., Kim, V.N. and Chang, H. (2020) The Architecture of SARS-CoV-2 Transcriptome. Cell, 181, 914–921 e910.

47. Galan, C., Sola, I., Nogales, A., Thomas, B., Akoulitchev, A., Enjuanes, L. and Almazan, F. (2009) Host cell proteins interacting with the 3’ end of TGEV coronavirus genome influence virus replication. Virology, 391, 304–314.

48. Wu, H.Y., Ke, T.Y., Liao, W.Y. and Chang, N.Y. (2013) Regulation of coronaviral poly(A) tail length during infection. PLoS One, 8, e70548.

49. Kawahara, H., Imai, T., Imataka, H., Tsujimoto, M., Matsumoto, K. and Okano, H. (2008) Neural RNA-binding protein Musashi1 inhibits translation initiation by competing with eIF4G for PABP. J Cell Biol, 181, 639–653.

50. Chen, C., Nadeau, S., Yared, M., Voinov, P., Xie, N., Roemer, C. and Stadler, T. (2022) CoV-Spectrum: analysis of globally shared SARS-CoV-2 data to identify and characterize new variants. Bioinformatics, 38, 1735–1737.

51. Conde, L., Allatif, O., Ohlmann, T. and de Breyne, S. (2022) Translation of SARS-CoV-2 gRNA Is Extremely Efficient and Competitive despite a High Degree of Secondary Structures and the Presence of an uORF. Viruses, 14.

52. Chan, A.P., Choi, Y. and Schork, N.J. (2020) Conserved Genomic Terminals of SARS-CoV-2 as Coevolving Functional Elements and Potential Therapeutic Targets. mSphere, 5.

53. Chen, Z., Wang, C., Feng, X., Nie, L., Tang, M., Zhang, H., Xiong, Y., Swisher, S.K., Srivastava, M. and Chen, J. (2021) Interactomes of SARS-CoV-2 and human coronaviruses reveal host factors potentially affecting pathogenesis. EMBO J, 40, e107776.

54. Gordon, D.E., Jang, G.M., Bouhaddou, M., Xu, J., Obernier, K., White, K.M., O’Meara, M.J., Rezelj, V.V., Guo, J.Z., Swaney, D.L. et al. (2020) A SARS-CoV-2 protein interaction map reveals targets for drug repurposing. Nature, 583, 459–468.

55. Ramani, A., Muller, L., Ostermann, P.N., Gabriel, E., Abida-Islam, P., Muller-Schiffmann, A., Mariappan, A., Goureau, O., Gruell, H., Walker, A. et al. (2020) SARS-CoV-2 targets neurons of 3D human brain organoids. EMBO J, 39, e106230.

56. Lin, J.Y., Kuo, R.L. and Huang, H.I. (2019) Activation of type I interferon antiviral response in human neural stem cells. Stem Cell Res Ther, 10, 387.

57. Wu, X., Dao Thi, V.L., Huang, Y., Billerbeck, E., Saha, D., Hoffmann, H.H., Wang, Y., Silva, L.A.V., Sarbanes, S., Sun, T. et al. (2018) Intrinsic Immunity Shapes Viral Resistance of Stem Cells. Cell, 172, 423–438 e425.

58. Wu, X., Kwong, A.C. and Rice, C.M. (2019) Antiviral resistance of stem cells. Curr Opin Immunol, 56, 50–59.

59. Dauber, B. and Wolff, T. (2009) Activation of the Antiviral Kinase PKR and Viral Countermeasures. Viruses, 1, 523–544.

60. Kim, Y., Lee, J.H., Park, J.E., Cho, J., Yi, H. and Kim, V.N. (2014) PKR is activated by cellular dsRNAs during mitosis and acts as a mitotic regulator. Genes Dev, 28, 1310–1322.

61. Lu, Z. and Hunter, T. (2009) Degradation of activated protein kinases by ubiquitination. Annu Rev Biochem, 78, 435–475.

