## Supplemental figures for "RNA binding protein Musashi1 interacts with the viral genomic RNA and restricts SARS-CoV-2 infection by repressing translation" (from another version of this preprint: version 1, 2024-10-01)

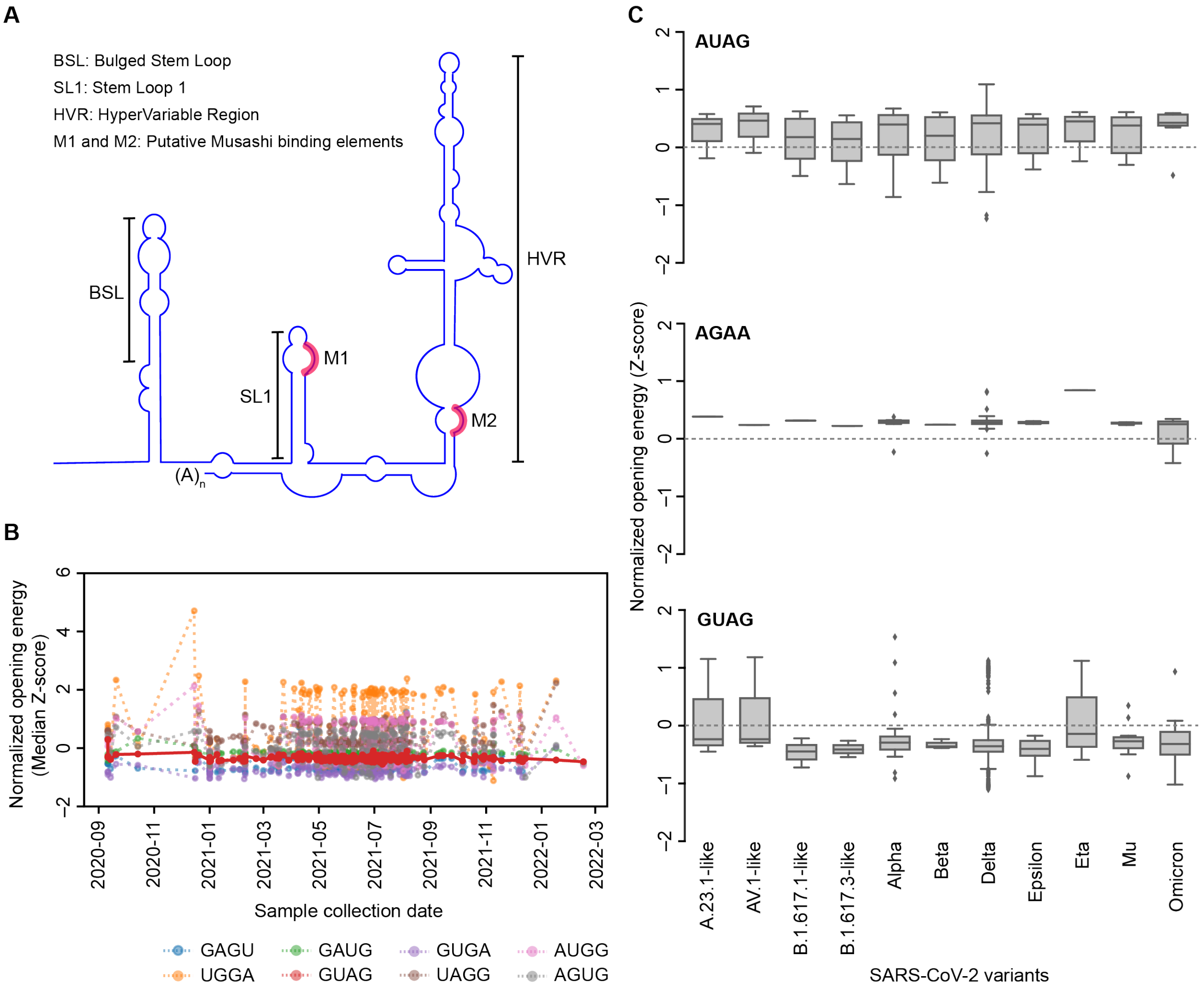


**Supplementary Figure 1. Musashi Binding Elements in the 3'UTR of SARS-CoV-2 genome.**

**(A)** The 3'UTR of SARS-CoV-2 forms a distinct secondary structure with two prominent Musashi Binding Elements (MBEs)- M1 and M2 in the stem-loop1 (SL1) and hyper variable region (HVR), respectively. **(B)** Graph showing the Z-scores of the normalised opening energy of different motifs in SARS-CoV-2 variants, with randomized GUAG consensus, collected over different time points. **(C)** Boxplot of distributions of the Z-scores of the normalised opening energy of the different consensus MBEs (AUAG, upper panel; AGAA, middle panel and GUAG, lower panel) among different SARS-CoV-2 variants.

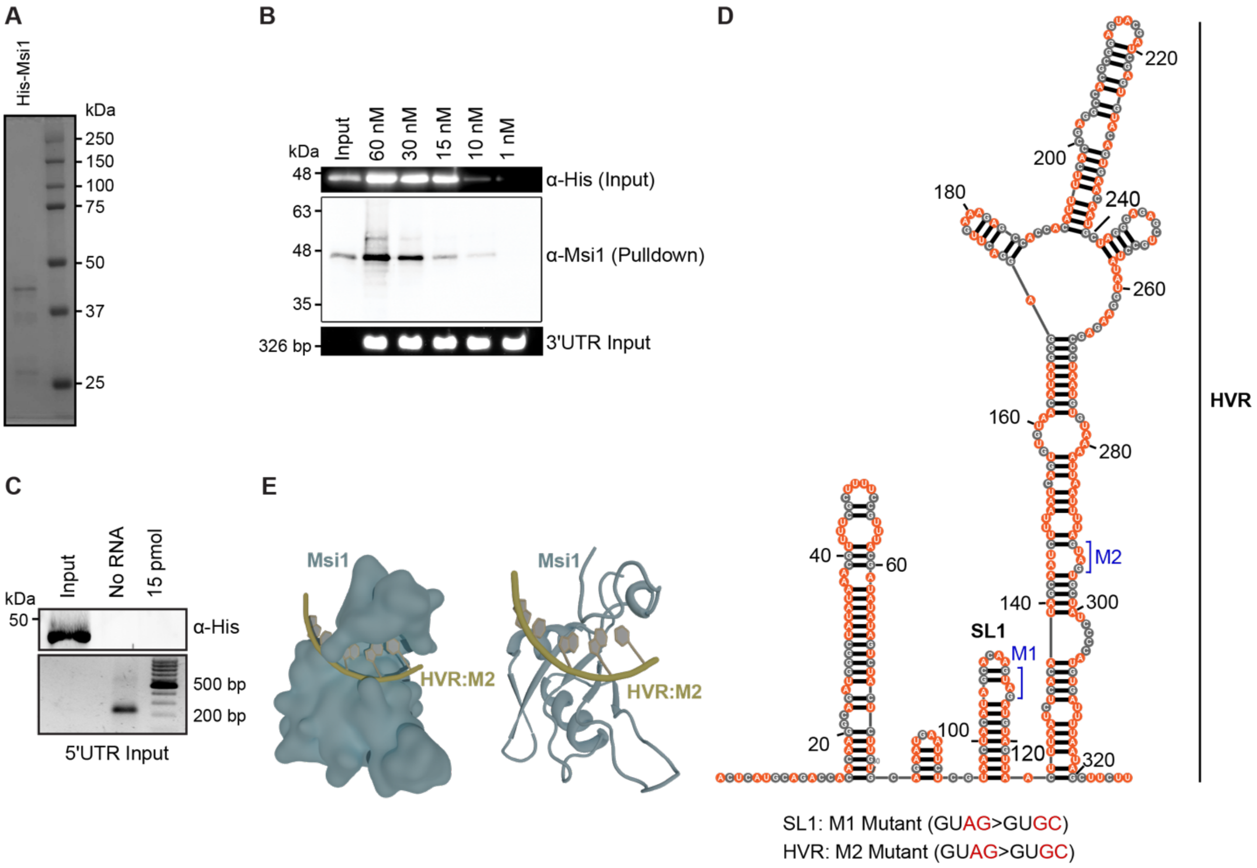


**Supplementary Figure 2. Msi1 directly interacts with SARS CoV2 3'UTR via MBEs.**

**(A)** Commassie gel showing purified His tagged Msi1 protein **(B)** RNA pull-down of different concentrations of purified Msi1 with a constant concentration of biotinylated 3'UTR of SARS-CoV-2 (15 pmol), followed by immunoblot analysis with His tag antibody. The upper panel shows Msi1 protein input, and the lower panel shows RNA input. **(C)** RNA pull-down of purified Msi1 protein with *in vitro* transcribed and biotinylated 5'UTR of SARS-CoV-2, followed by immunoblot analysis probed with His-tag antibody. **(D)** Secondary structure of 3'UTR of SARS-CoV-2 indicating the specific mutations in motifs M1 and M2. **(E)** Multifaceted visualization of Msi1 (PDBID:5X3Z) and SARS-COV-2 M2 RNA motif interaction. The left panel shows the surface representation of the Msi1 in complex with viral motif M2, highlighting the detailed interaction surface, including key binding sites and molecular contours. The right panel presents a cartoon representation of the same complex, emphasizing the overall structural arrangement and folding patterns of the Msi1 and the viral motif M2.

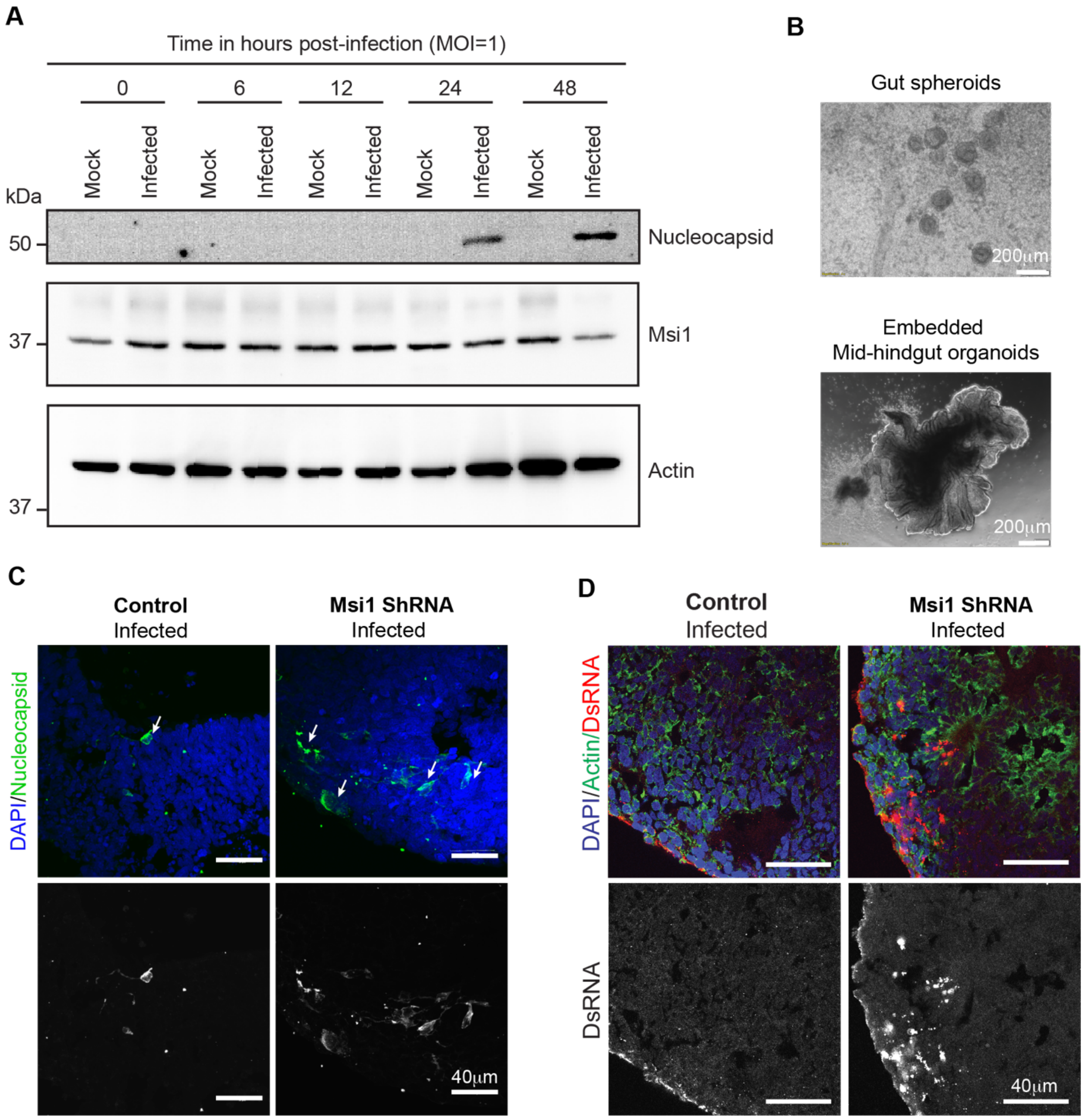


**Supplementary Figure 3. Msi1 depletion promotes viral replication in 3D cultures.**

**(A)** Time course of SARS-CoV-2 infection in Caco-2 cells showing that the levels of Msi1 are not affected. **(B)** Representative images of gut spheroids and mid-hindgut organoids used for SARS-CoV-2 infection. Immunostaining of **(C)** Nucleocapsid (green) and **(D)** DsRNA (red) and Actin (green) in organoids treated with control or Msi1 ShRNA (MOI1) after 48 h. DAPI is stained in blue for both.

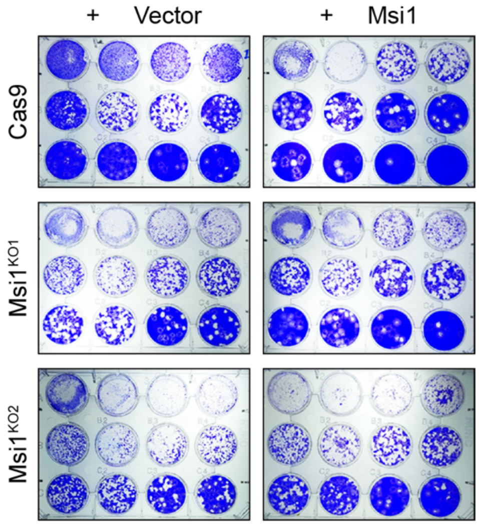


**Supplementary Figure 4. Msi1 represses viral translation.**

Viral Plaque assay depicts the difference in plaque formation between Wildtype Caco-2 and Msi1 KO cell line with transgene expression of Msi1 or empty vector.

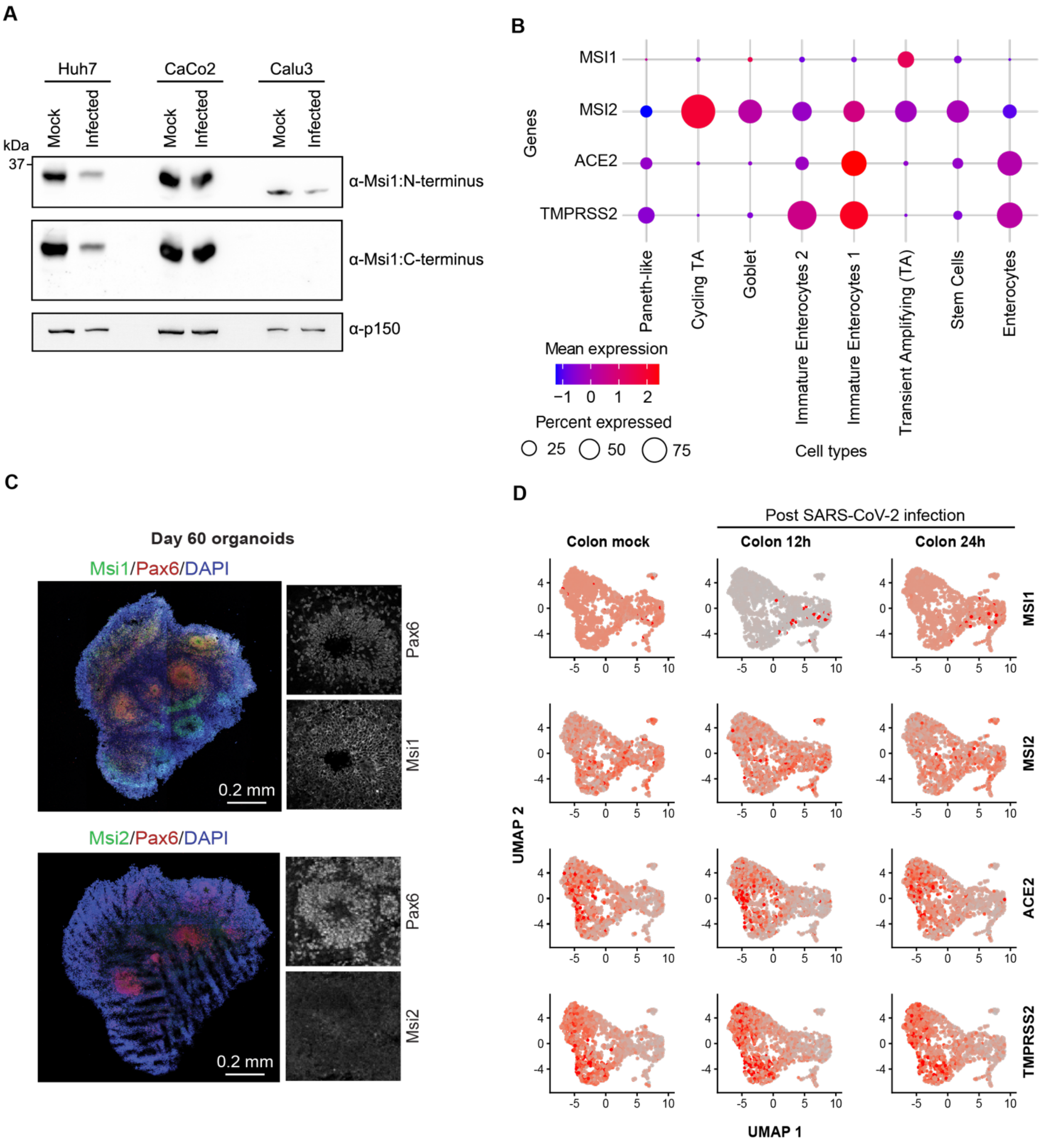


**Supplementary Figure 5. Msi1 expression upon SARS-CoV-2 infection.**

**(A)** Western blot analysis of Msi1 in Huh7, Caco-2 and Calu3 cell lines after mock or SARS-CoV-2 infection. P150 was used as a loading control. **(B)** Bubble plot, depicting expression levels of MSI1, ACE2 and TNPRSS2 across cell clusters in the intestine from publicly available dataset (1). The color of each bubble represents mean level of expression of the indicated gene whereas the relative size of the bubble is indicative of the percentage of cells expressing the gene. **(C)** Confocal image analysis of a D60 cerebral brain organoid expressing Msi1(green) and Pax6 (red) or Msi2 (green) and Pax6 (red). DAPI (blue) is used to stain nuclei. **(D)** Uniform manifold approximation and projection (UMAP) embedding of the scRNAseq data of infected colon organoids at 12 and 24 hours post infection, colored by the corrected targeted normalized expression of SARS-CoV-2.

**Supplementary References**

1. Triana, S., Metz-Zumaran, C., Ramirez, C., Kee, C., Doldan, P., Shahraz, M., Schraivogel, D., Gschwind, A.R., Sharma, A.K., Steinmetz, L.M. *et al.* (2021) Single-cell analyses reveal SARS-CoV-2 interference with intrinsic immune response in the human gut. *Mol Syst Biol*, **17**, e10232.
